# Mgat4b mediated selective *N*-glycosylation regulates melanocyte development and melanoma progression

**DOI:** 10.1101/2024.10.10.617552

**Authors:** Babita Sharma, Keerthic Aswin, Tanya Jain, Ayesha Nasreen, Ayush Aggarwal, Yogaspoorthi Subramaniam, Jeyashri Rengaraju, Srashti Jyoti Agrawal, Mayank Bhatt, Bhaskar Paul, Koushika Chandrasekaran, Aanchal Yadav, Jyoti Soni, Rajat Ujjainiya, Md Quasid Akhter, Shantanu Sen Gupta, Rajesh Pandey, Shruthy Suresh, Srinivasa-Gopalan Sampathkumar, Vivek T Natarajan

**Affiliations:** CSIR-Institute of Genomics and Integrative Biology, Mathura Road, New Delhi 110025, India; Academy of Scientific and Innovative Research (AcSIR), Ghaziabad, Uttar Pradesh 201002, India; Laboratory of Chemical Glycobiology, National Institute of Immunology, Aruna Asaf Ali Marg, New Delhi, 110067, India

## Abstract

Melanocyte development involves key pathways that are often recapitulated during melanoma initiation, highlighting the importance of understanding the regulators that control these early processes and also contribute to cancer onset. Our study identifies *mgat4b*, a glycosyl transferase involved in selective *N*-glycan branching enriched in pigment progenitors, as a key regulator of directional melanocyte migration and establishment of melanocyte stem cell (McSC) pool during early development. Single cell RNA (scRNA) sequencing analysis in zebrafish upon targeted disruption of *mgat4b* reveals, that migratory melanocyte progenitors marked by galectin expression fail to persist. Lectin affinity proteomic analysis reveals the glycosylation of key melanocyte proteins GPNMB, KIT, and TYRP1 to be under the control of MGAT4B in melanocytic cells. Additionally, mislocalization of Junctional plakoglobin (JUP) explains the observed defects in cell adhesion and migration to be regulated by MGAT4B but not its isozyme MGAT4A. Our meta-analysis further reveals that melanoma patients with both the BRAF^V600E^ mutation and elevated MGAT4B levels have significantly worse survival outcomes compared to those with only the BRAF^V600E^ mutation. By leveraging the MAZERATI platform to model BRAF^V600E^ driver mutation *in vivo*, we show that *mgat4b* mutant cells fail to aggregate and initiate tumors. RNA profiling of the transformed melanocytes revealed cell-cell junction, adhesion and ECM binding to be probable contributing factors that resulted in the failure of tumor onset. Using a small-molecule inhibitor we demonstrate the inhibitory role of this complex *N*-glycosylation in the progression of early-stage melanoma. Our study underscores the importance of selective *N*-glycan branching in both melanocyte development and melanoma initiation, suggesting MGAT4B as a promising therapeutic target for melanoma treatment.

## Introduction

Melanocytes, are specialized cells that originate from neural crest cells (NCCs) and containing melanin-rich melanosomes. These cells are crucial for skin pigmentation that serves a protective function against DNA damage (Mort et al., 2015). Early specifying melanocytes directly originate from the neural crest and migrate over the developing vertebrate embryo to orchestrate the pigmentation pattern (Brenner & Hearing, 2008; Brombin & Patton, 2024). During regeneration, these cells are replenished by tissue resident Melanocyte stem cell (McSC) pool (Brombin et al., 2022). Unravelling the molecular mechanisms of melanocyte development is vital not only for understanding early development but also for combating diseases like melanoma, as these developmental processes are often co-opted during the transformation of melanocytes to melanoma and its further progression (Baggiolini et al., 2021; Kaufman et al., 2016).

Recent research has elucidated important pathways such as Kit and ErbB signalling involved in establishing McSCs (Brombin et al., 2022; Dooley et al., 2013; Kelsh & Barsh, 2011; Minchin & Hughes, 2008). Melanocytes originate directly from NCCs and from multipotent progenitor cells that reside in tissues but are derived from NCCs during early development (Brombin & Patton, 2024). The transcription factor Mitfa plays a central role in this process, with additional support from factors like tfap2a-c (Seberg et al., 2017). Thyroid hormone influences terminal differentiation, limiting the final numbers of mature melanocytes (Saunders et al., 2019). Adult pigment cell stripes are formed by the Schwann cell precursors providing a secondary source of melanocytes, marked by a distinct gene expression pattern (Adameyko & Lallemend, 2010; Adameyko et al., 2009; Dupin et al., 2003). Furthermore, recent studies reveal an important role for Prl3-Ddx21 axis, in preserving McSC pools (Johansson et al., 2020). Interestingly these factors also have a central role to play in melanoma cells, highlighting the recapitulation of developmental programs in melanoma (Johansson et al., 2020; Lal et al., 2013; Pham et al., 2020; Tiwary et al., 2014).

There is growing evidence that epigenetic and transcriptional developmental programs specific to melanocytes and their precursors are re-expressed during melanoma initiation (Baggiolini et al., 2021; Kaufman et al., 2016). It is important to identify the programs regulating melanocyte cell states as this may help us identify and target early events in melanoma initiation. In this study, we set out to identify factors that promote melanocyte differentiation from neural crest cells and found that a glycosyl transferase *mgat4b* initiates the formation of a unique *N*-glycan branch on set of glycoproteins, and functions as a crucial orchestrator of melanocyte development.

We report that *mgat4b* as a selective *N*-glycosyl modifier is enriched in pigment progenitors and melanophores during early melanocyte development and demonstrate that it is crucial for melanocyte migration and patterning in zebrafish *in vivo* as well as in cultured mammalian melanocytes. Melanocyte specific targeting of *mgat4b* abrogates migratory cells marked by Galectin1 expression. Differential lectin affinity proteomics revealed that the transmembrane protein GPNMB, the receptor tyrosine kinase KIT and TYRP1 are modified by MGAT4B. Additionally, expression levels and localization of a sub-membranous protein Gamma catenin (JUP) is altered in the mutant group, contributing to the aberrant cell adhesion and migration defects observed in these animals. From publicly available data-sets we observe that MGAT4B is overexpressed in human melanoma. Using MAZERATI, a platform designed to swiftly model potential cancer drivers *in vivo* (Ablain et al., 2018), we demonstrate that *mgat4b* is necessary for melanoma initiation. Thus, our work highlights the role of selective *N*-glycosylation in early-stage melanocyte development and how this impacts melanoma initiation. These findings underscore the importance of selective *N*-glycan branching in melanoma development and positions MGAT4B as a promising target for melanoma therapy.

## Results

### *mgat4b* is enriched in pigment progenitors and melanophores

In an anticipation of finding factors that favours melanocyte development from NCCs, we performed re-analysis of zebrafish scRNA data in neural crest cells (Saunders et al., 2019). Dimensionality reduction and unsupervised clustering was applied to the entire population of sox10+ cells, leading to the identification of markers specific to pigment progenitors and developing melanocytes (Figure 1A). As expected, *miGa*, *Gap2a* and *mlpha* were enriched in melanophore and pigment progenitors (Figure 1B). Interestingly, we observed that *mgat4b*, a gene encoding a glycosyl transferase, was preferentially enriched in melanophores compared to other chromatophores which arise from the same cell population of pigment progenitors (Figure 1B). Mgat4b expression was higher in the progenitors compared to mature melanophores suggesting an early role, if any (Figure 1C). Mgat4b is an enzyme located in golgi-apparatus that forms β1→4 linked branch on α1→3 mannose residues of selective *N-* glycoproteins (Figure 1D). In this subpopulation of *mgat4b*-expressing cells, receptor-ligand analysis highlighted key factors involved in cell migration, such as Plexin b2 and Nectin2, which are crucial for migration-related signalling. Additionally, the presence of Integrin α4, α5, and α1, along with ADAM10, is suggestive of their role in interactions with the extracellular matrix essential for effective migration (Figure 1E) (Gu & Taniguchi, 2008). Since many of the predicted targets of *mgat4b* modulates cell surface receptor signalling and governs cell-cell or cell-matrix adhesion, we reasoned that *mgat4b* may play a critical role in melanocyte patterning and development.

**Figure 1:**
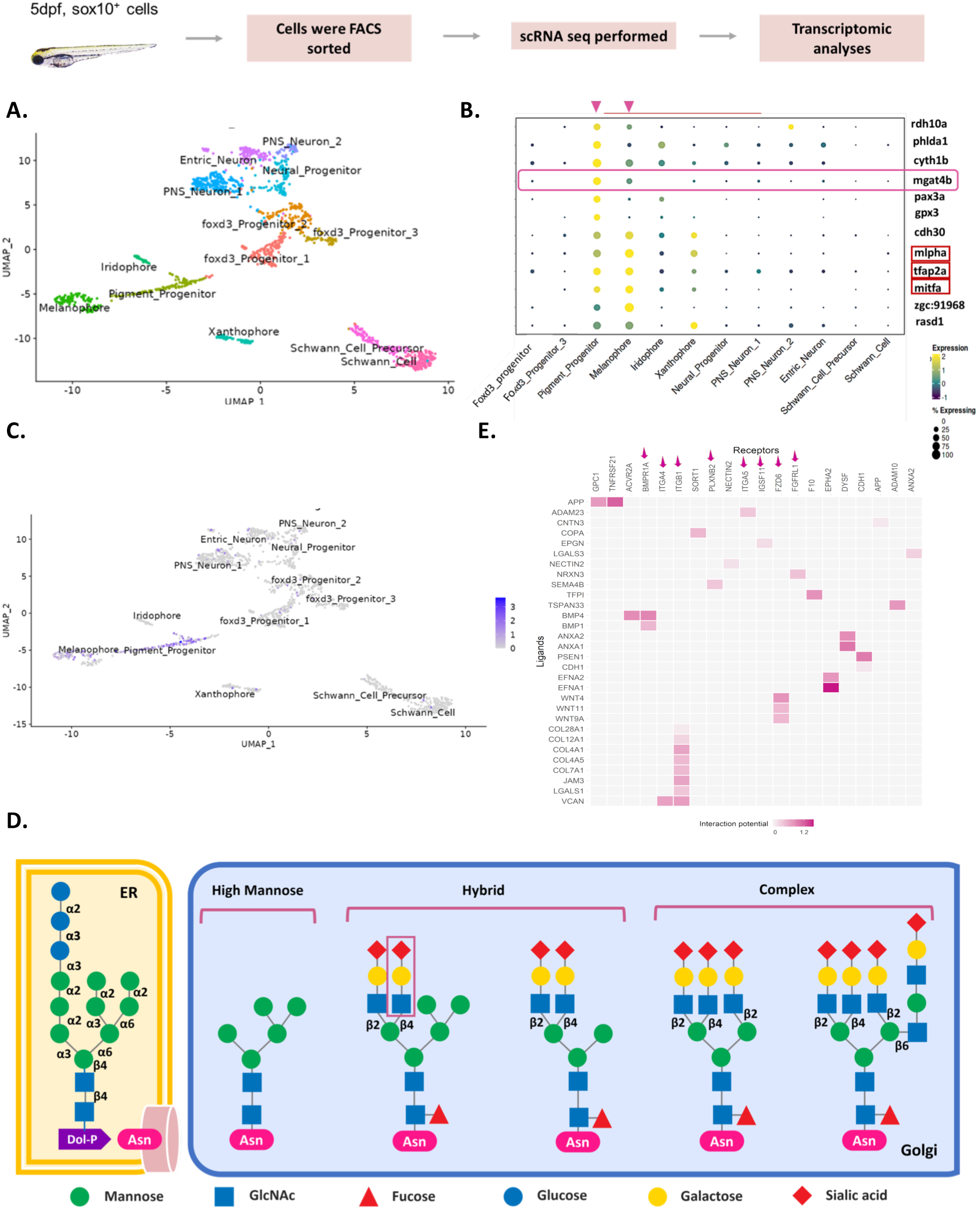
mgat4b is enriched in Pigment progenitors and Melanophores: **A.** UMAP visualization of the Zebrafish Sox10+ve cells colored by the identified states **B.** Dot plot showing the top marker genes enriched in each cluster with the size showing the percent of cell expressing the gene and color showing the scaled mean expression value in each cluster (Wilcoxon-Mann-Whitney test with average log fold change > 0.25 and adjusted p value ≤ 0.05). List includes known cell-type marker genes (marked in red box) and new candidate markers **C.** UMAP plot shows enrichment of *mgat4b* in pigment progenitor and melanophore clusters **D.** Mammalian protein *N*-glycosylation starts in the endoplasmic reticulum and continues in the Golgi apparatus, where three mature types of *N*-glycan structures—high-mannose, hybrid, and complex—are formed. The enzyme GT-4b transfers *N*-acetylglucosamine in a β1→4 linkage to α1→3-linked mannose, which kickstarts the creation of a specific N-glycan branch. This branch often includes a sialylated *N*-acetyllactosamine sequence, highlighting its distinctive features in glycan structure **E.** Heatmap of NicheNetR-identified ligand/receptor pairs indicating interaction potential between *mgat4b*+ve receiver cells and other sender cells sampled by scRNAseq of zebrafish Sox10+ve cells.

### Global targeting of *mgat4b* results in melanophore patterning defects

To understand the role of *mgat4b* in melanocyte development, we adopted three independent strategies. First, we used a splice block morpholino against *mgat4b* to transiently knockdown the gene. We observed mild head and eye size reduction in the morphants (Supplementary Fig 1A, B). We analysed these animals till day 5, when the melanophores are well patterned, melanized and all five melanophore stripes dorsal, ventral, yolk sac, and two lateral are visible. *mgat4b* morphants showed a significantly lower number of lateral mid-line melanophores with respect to control animals (Supplementary Fig 1C, D).

To assess the initial melanophore emergence along the crest, we utilized a transgenic zebrafish line Tg(Mitfa:GFP). Fluorescence imaging conducted at 24hpf showed reduction in emergent streams of Mitfa+cells from the crest in the absence of *mgat4b* (Supplementary Figure 1 E, F). Furthermore, using a Tg(ftyrp:GFP) line where GFP expression is specific to differentiating melanophores, we found aberrant melanophores patterning in the trunk region upon *mgat4b* knockdown at 2dpf (Supplementary Figure 1 G, H).

Secondly, to complement the morpholino approach, we generated CRISPR mediated global knockout of *mgat4b in vivo* by injecting the sgRNA-Cas9 RNP complex at the single-cell stage (Supplementary Figure 2A). Brightfield imaging of probable F0 mutants (M4b mut) and non-targeting control embryos at 3dpf revealed patterning defects, with a notable portion of melanophores becoming arrested along the path (Supplementary Figure B-D). Sequencing confirmed an in-del mutation disrupting the coding sequencing of *mgat4b* in F1 (Supplementary Figure 2E). The mutant embryos exhibited a decreased number of lateral melanophores compared to the WT animals (Supplementary Figure 2D). Thus, the global mutant of *mgat4b* recapitulated the morpholino phenotype, with a decrease in the number of lateral melanophores and overall patterning defects.

### Melanocyte specific ablation of *mgat4b* leads to patterning abnormalities

Thirdly, in order to assess if the defects observed were due to melanocyte specific functions of *mgat4b*, we adopted a cell-type specific mutation strategy by modifying the MinicoopR-sgRNA construct developed for rescuing melanophores in *casper* animals. We replaced *miGa* ORF with a visualization marker GFP, so that melanocyte specific expression of this construct could be visualized through co-expression of GFP with a concomitant ability to target individual genes in melanophores (Supplementary Figure 3A). We injected this Mitfa: Cas9, Mitfa: gfp plasmid at the single cell stage and observed majority of melanophores labelled at 2dpf (Supplementary Figure 3A). This construct was specifically expressed in melanocytes as GFP+ sorted cells showed enrichment for melanocyte-specific markers such as *miGa, dct,* and *tyr* at 2 dpf (Supplementary Fig 3B). Whole mount in-situ (WISH) was used to check for Cas9 RNA expression, which was well-restricted to areas where melanophores reside based on the position of the GFP signal (Supplementary Figure 3C) (Raja, Subramaniam, et al., 2020).

We cloned *mgat4b* sgRNAs into this Mitfa:Cas9;Mitfa:gfp and specifically ablated *mgat4b* in melanophores (Supplementary Figure 3 E). This revealed patterning defects at the 3-day post-fertilization (dpf) stage (Supplementary Figure 3 F, H). By 5 dpf, there was a notable decrease in the number of lateral melanophores, (Supplementary Figure 3 G, I). Thus, all three experimental approaches that were employed to target *mgat4b* consistently showed a reduction in the number of embryonic lateral-line melanophores. Thereby, our somatic cell-type ablation strategy highlights that gene ablation in melanocyte specific manner *in vivo* is a powerful tool for screening candidates and evaluating their roles using pigmentation outcomes as a readout. This strategy highlights the melanocyte-specific role of *mgat4b*, demonstrated by the mutant’s inability to form lateral melanophores. To unravel the mechanistic basis of this phenotype, we utilized single-cell RNA sequencing in GFP-positive cells.

### scRNA data maps the fate of *mgat4b* targeted melanocyte precursors

Our data thus far demonstrates that *mgat4b* mutant results in dysregulation of melanophore development. However, *mgat4b* loss seems to specifically impact lateral melanophores more than the dorso-ventral ones. Thus, to identify the key melanophore populations altered by *mgat4b*, we performed single-cell RNA sequencing (scRNA-seq) on control and mutant GFP+ cells at 36 hpf using the 10X Chromium platform. After integrating all samples, we identified six distinct clusters (Figure 2A, B). Based on markers from previously published datasets (Brombin et al., 2022; Saunders et al., 2019), we assigned the identities for these clusters. We identified two proliferative neural crest (NC) populations: twist1a+ NC progenitors and foxd+ progenitors (Figure 2C). Additionally, we identified three distinct clusters expressing combination of different chromatophore markers. One cluster expressed markers of all three types of chromatophores namely xanthophore, melanophore and iridophore (expressing aox5, mitfa, pnp4a), stem cell markers (crestin, sox10) and migratory markers (rac2, csf3b, rgs2 and itga6) (Figure 2C and Supplementary Figure 4D). Hence, we designated this cluster as the migratory MIX+ population. We observed this cluster to be also be enriched for tyrp1b and not foxd3 (Figure 2C), therefore, it is likely that they will eventually become mature melanophores (Curran et al., 2010). In addition, we identified two bipotent clusters, MX+ and MI+. Representative markers for each cluster are shown in the dot plot and UMAP (Figure 2B, C) and top ten markers of each cluster are shown as heatmap (Sup Figure 4C). Importantly, the migratory MIX+ cluster was specifically depleted in *mgat4b* mutant group suggesting that these cells may contribute to the melanocyte defects observed *in vivo* (Fig 2D&E). Further analysis revealed that lectin genes such as *lgals1, lgals3 and lgals9* are highly enriched in the MIX+ cluster (Supplementary Figure 4D). These lectin genes have galactoside-binding properties, and are known to interact with substrates that contain branched structures modified by Mgat4 (Varki et al., 2015). This suggests a critical requirement of specific *N*-glycosylation branching mediated by *mgat4b* for the sustenance of this migratory population.

**Figure 2:**
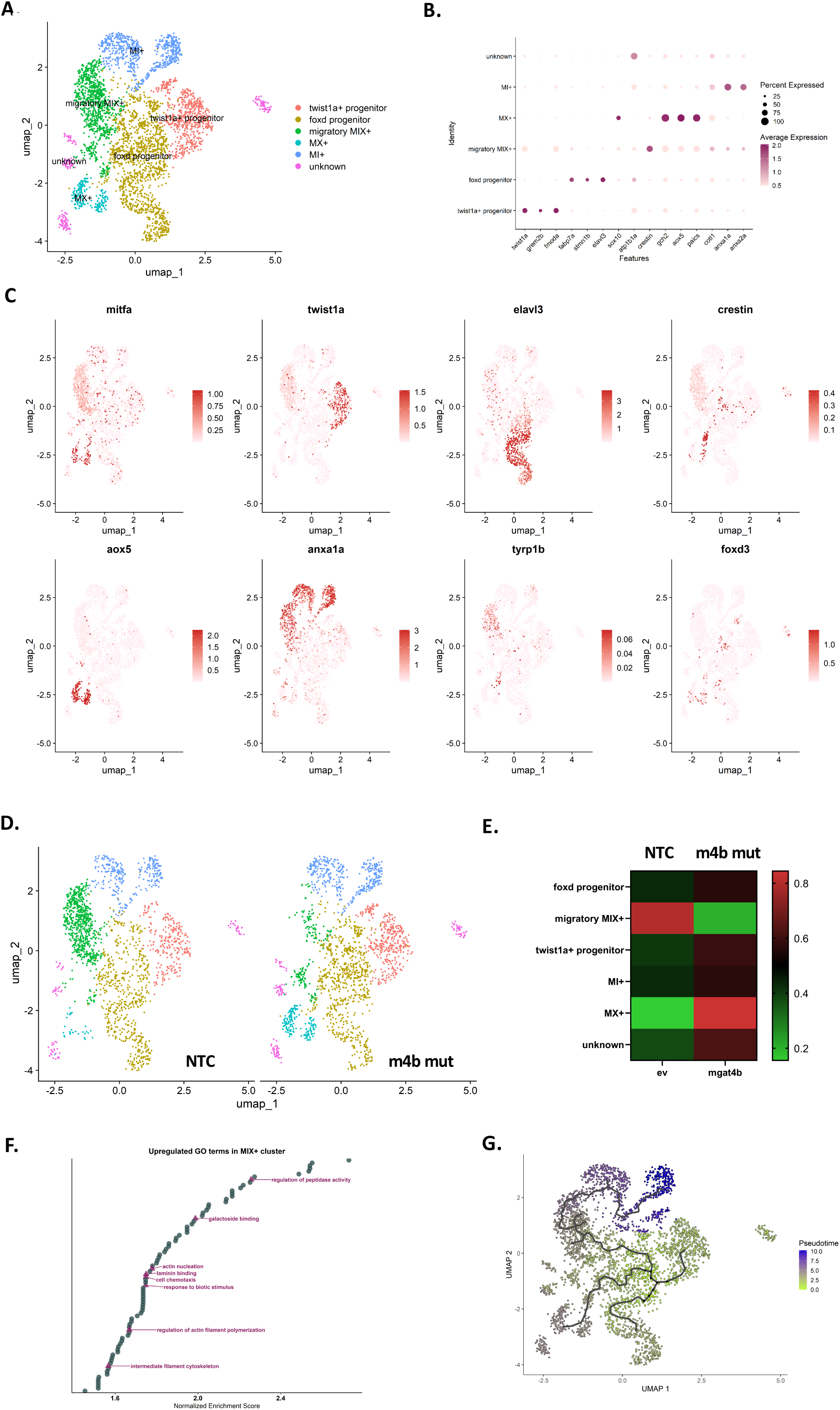
Melanocyte specific ablation of *mgat4b* leads to cell-arrest and results in cell death: **A.** UMAP visualization of the zebrafish Mitfa+ve cells integrated from Non-targeted control (NTC) and melanocyte specific knock out of *mgat4b* (m4b mut)cells colored by the identified states **B**. Dot plot showing the top marker genes enriched in each cluster with the size showing the percent of cell expressing the gene and color showing the scaled mean expression value in each cluster **C**. UMAPs of Mitfa+ cells with color change from light pink (negative) to red based on log normalized scaled expression of mitfa, twist1a, elavl3, crestin, aox5, anxa1a and tyrp1b and foxd3 **D**. UMAP visualization of the Zebrafish Mitfa+ve cells from NTC and m4b mut cells colored by the identified states **E**. Heat map showing the proportion of cells belonging to certain cell-type distributed between NTC and m4b mut samples **F**. Waterfall plot depicting top upregulated GO-terms in MIX+ cluster plotted against their normalized enrichment score **G**. Pseudotime ordering of the cells, coloring based on pseudotime scores (Wilcoxon-Mann-Whitney test with average log fold change > 1 and adjusted p value ≤ 0.05)

Functional enrichment analysis of genes in the missing cluster identified significant enrichment of ECM binding, cell adhesion, and galectin binding (Figure 2F). We then performed cell-cell communication analysis to understand the cellular interactions in this cluster (Supplementary Figure 4A, B). The results confirmed our earlier observation of their migratory potential, as these cells expressed key receptors and metalloproteases (ITGA6, RAC2, MMP9, CXCL14 and MMP30) essential for cell migration, adhesion and chemotaxis-related processes (Supplementary Figure 4D). To investigate the lineage relationships between clusters, we conducted a pseudo-time analysis based on gene expression that revealed a trajectory. This revealed a branch point originating from NCs that transitioned into MX+ and MI+ through migratory MIX+ cells (Figure 2G). The specific reduction in the MIX+ cluster caused by the *mgat4b* mutation suggests an effect on melanoblast formation, while the MX+ and MI+ clusters remain unaffected. These are likely to differentiate into xanthophores and iridophores, as indicated by the expression of markers like aox5 for xanthophores and anxa1a and anxa2a for iridophores (Fig B, E and G). Consistent with this, there is no reduction in the xanthophore and iridophore populations in *mgat4b* mutant group. To summarize, through single cell transcriptomics, we identified that selective *N*-glycan branching mediated by *mgat4b* in migratory melanophore precursors is critical and essential for the early migration, directionality, and survival of cells that develop into melanophores.

### Mgat4b targeting leads to migration arrest and subsequent cell death

The NCC derived progenitors migrate dorso-medially to establish the McSC pool (Dooley et al., 2013). We therefore traced the dorso-medial migration of NCC derived progenitors. Live imaging of control animals showed directional migration of melanophores, with elongated dendrites extending in the direction of movement (Figure 3A, Supplementary Video 1). In contrast, *mgat4b* mutant animal exhibited melanophores extending dendrites in all directions. These melanophores appeared to be arrested along their paths and showed very limited movement (Figure 3B, Supplementary Video 2). Thus, underscoring the importance of *mgat4b* in facilitating the migration of melanocytes along their paths while maintaining the correct directional orientation.

**Figure 3:**
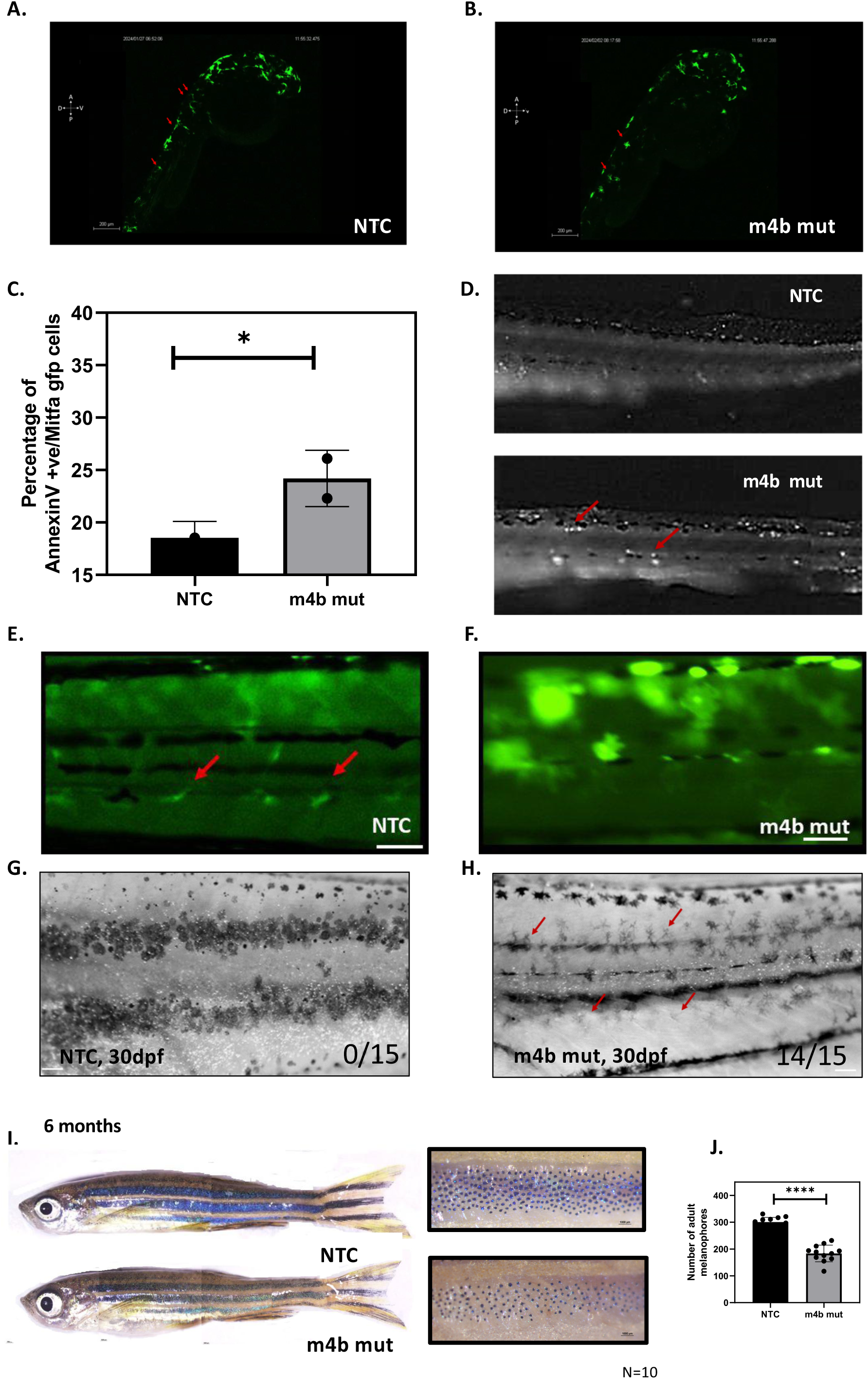
Melanocyte specific ablation of *mgat4B* leads patterning abnormalities and reduced adult melanophores. **A**, **B**. Snapshot of timelapse imaging showing early melanophore migration. Red arrows in NTC animal shows elongated melanophores migrating in dorsolateral or dorsomedial direction whereas in m4b mut animal they mark direction-less star-shaped melanophores, Scale bar-200µm **C**. Bar graph represents mean±s.e.m. of percentage of Annexin V+/Mitfa gfp+ cells in NTC and m4b mut animals **D**. Lateral images of 2dpf NTC and m4b mutant animal showing acridine orange puncta. Red arrows show puncta in the region where melanophore residing regions, Scale bars: 50 μm **E, F.** Lateral view of tissue specific NTC and m4b mut at 7dpf showing melanocyte stem cells (McSCs) residing pattern, Red arrows show the arrangement of McSCs in the trunk region, Scale bar: 50 µm **G, H**. Lateral view of 1 months old NTC and m4b mut animal showing pigment stripes after metamorphosis, Scale bar: 50 µm **I.** Lateral view of 6 months old NTC and m4b mut animal showing adult pigment stripes, the zoomed images on the right shows constricted melanophores upon epinephrine treatment, Scale bar: 1000 µm **J**. Bar graph represents mean±s.e.m. of melanophore count in adult stripes in NTC and m4b mut animals *P ≤ 0.05, **P ≤ 0.01, ***P ≤ 0.001, ****P ≤ 0.0001 and ns P > 0.05

To explore the fate of these arrested cells, we assessed the survival of these cells by Annexin V staining. A significant and consistent increase (from 18 % to 26%, p=0.0138) in the early apoptotic cell population was observed in *mgat4b* mutant animals compared to controls (Figure 3C). Furthermore, acridine orange staining was performed to examine the spatial distribution of these apoptotic cells. We observed a markedly higher frequency of GFP puncta in the trunk region of *mgat4b* mutant animals compared to control (Figure 3D). This region corresponds to where melanophores were arrested during their migration (Figure 3B). Hence it is likely that the failure to achieve the appropriate spatial positioning and an independent, or a consequent loss of growth factor signalling within this critical time window may preclude the migratory cells from establishing the McSC pool. Since *mgat4b* mutants showed decrease in both embryonic lateral and adult melanophores, we hypothesized that the McSC pools is altered in these animals. Lateral and adult melanophores arise from an McSC pool that resides in dorsal root ganglion (DRG) (Mort et al., 2015). We set out to identify McSCs in 4 dpf animals as previously reported (Brombin et al., 2022; Dooley et al., 2013; O’Reilly-Pol & Johnson, 2013), and observed multiple dendritic Mitfa:gfp cells in the *mgat4b* mutant animals, in place of the few elongated McSCs associated with the DRG observed in the control animals (Figure 3E, F). These defects extended into one month old animals in which the stripe establishment is incomplete, the mutants display dendritic melanophores and are unable to orchestrate into the stripes (Figure 3 G, H). In the 6-month-old adult animals, while the stripes could be observed the melanonophore numbers were diminished in *mgat4b* mutant group compared to the NTC group (Figure 3 I, J). Therefore, *mgat4b* appears to have a role in establishing the McSC pool. It likely influences the MIX+ cells or their precursors during the migration phase of melanocyte development. However, the exact molecular mechanism behind this process remains unknown thus far.

### MGAT4B expression in melanocytes is governed by MITF

As *mgat4b* plays a key role in melanophore development, we assessed the changes in *mgat4b* expression in melanocytes across the developmental time-window. Reanalysis of zebrafish microarray data from our group showed that similar to the *miGa* expression, *mgat4b* expression peaks during the 34-36 hpf time-frame (Supplementary Figure 2F, GSE189059). This coincides with the migration and positioning of melanophores within embryonic stripe patterns in zebrafish development. Given that *miGa* is a key transcription factor responsible for these processes, we hypothesized that *mgat4b* may mediate its effects through a *miGa* program in melanocytes.

To test this, we first assessed if *miGa* overexpression could rescue defects observed in *mgat4b* mutant. We observed melanocyte patterning defects and reduced melanophore survival, even with increased *miGa* expression (Supplementary Figure 5 A-E). Suggesting that the effects of *mgat4b* are either independent of *miGa* or likely occur downstream of *miGa* in melanocytes. Indeed, studies have shown that glycosyltransferases are regulated by cell-type-specific transcription factors within precise spatio-temporal windows to perform specific functions (Ohtsubo et al., 2011). Thus, we reasoned that *miGa* could regulate *mgat4b* in melanocytes. To investigate this, we analyzed previously published single-cell RNA sequencing data obtained from zebrafish mitfa+ cells at 28 hpf and clustered them based on their levels of endogenous *miGa* expression (GSM7706846, GSM7706847). Strikingly, we observed a consistent correlation between the expression levels of *miGa* and *mgat4b* within these cells, suggesting that cells exhibiting high *miGa* expression also display elevated levels of *mgat4b* (Supplementary Figure 5F). To test if *miGa* transcriptionally regulates *mgat4b*, we overexpressed MITF in B16F10 mouse melanoma cells and assessed its effects on MGAT4B levels. MITF overexpression increased the protein levels of MGAT4B and decreased when MITF was downregulated (Supplementary Figure 4 G, H). Finally, we utilized ChIPqPCR to confirm the binding of MITF to the MGAT4B promoter region. This showed an enrichment of MITF in the MGAT4B promoter region (Supplementary Figure 5I). In conclusion, our findings demonstrate that MITFA transcriptionally regulates MGAT4B in melanocytes, playing a crucial role in melanophore migration, positioning, and survival.

### Loss of *Mgat4b* reduces invasiveness and disrupts the velocity and directionality of migrating melanocytes

To dissect the molecular mechanisms by which *Mgat4b* regulates melanocyte functions, we knocked out *Mgat4b* in B16 mouse melanoma cells (Supplementary Figure 6 A-D). The *Mgat4b* KO cells displayed distinct morphological differences compared to the control, including fewer cell-cell contacts and non-cohesive aggregate colonies (Figure 4A, B). Phalloidin staining of filamentous-actin revealed that the KO cells showed extended filopodia, whereas the control cells were more spindle-shaped and firmly adhered to the substratum (Figure 4C). Since functional enrichment analysis of zebrafish *mgat4b* mutants highlighted ECM binding, migration, and cell adhesion as the most enriched pathways, we evaluated the invasive capability of these *Mgat4b* KO cells using a Matrigel-coated trans-well invasion assay. We observed a significant decrease (p= <0.0001) in invading cells, reinforcing the hypothesis that *Mgat4b* could be involved in ECM degradation and extravasation (Figure 4D, E). To further explore this, we assessed the migration capability of KO cells using a chemotaxis assay. Wildtype and KO cells were seeded in a trough connected to a reservoir containing SCF (chemoattractant) on one side and control media on the other, creating a stable SCF gradient across the imaging field. We observed that wildtype cells preferentially migrated toward increasing SCF gradient and maintained persistent migration directionality throughout the imaging period (Figure 4F). In contrast, KO cells showed no preferential direction of migration and additionally moved with significantly lower velocity (Figure 4G, Supplementary table 1). This suggests that *Mgat4b* is crucial not only for the directional movement of melanocytes but the migration ability in itself. In this context, it is intriguing that the *Mgat4b* KO cells put out extended filopodial projections but are incompetent in cell migration (Fig 4G, Supplementary table 1).

**Figure 4:**
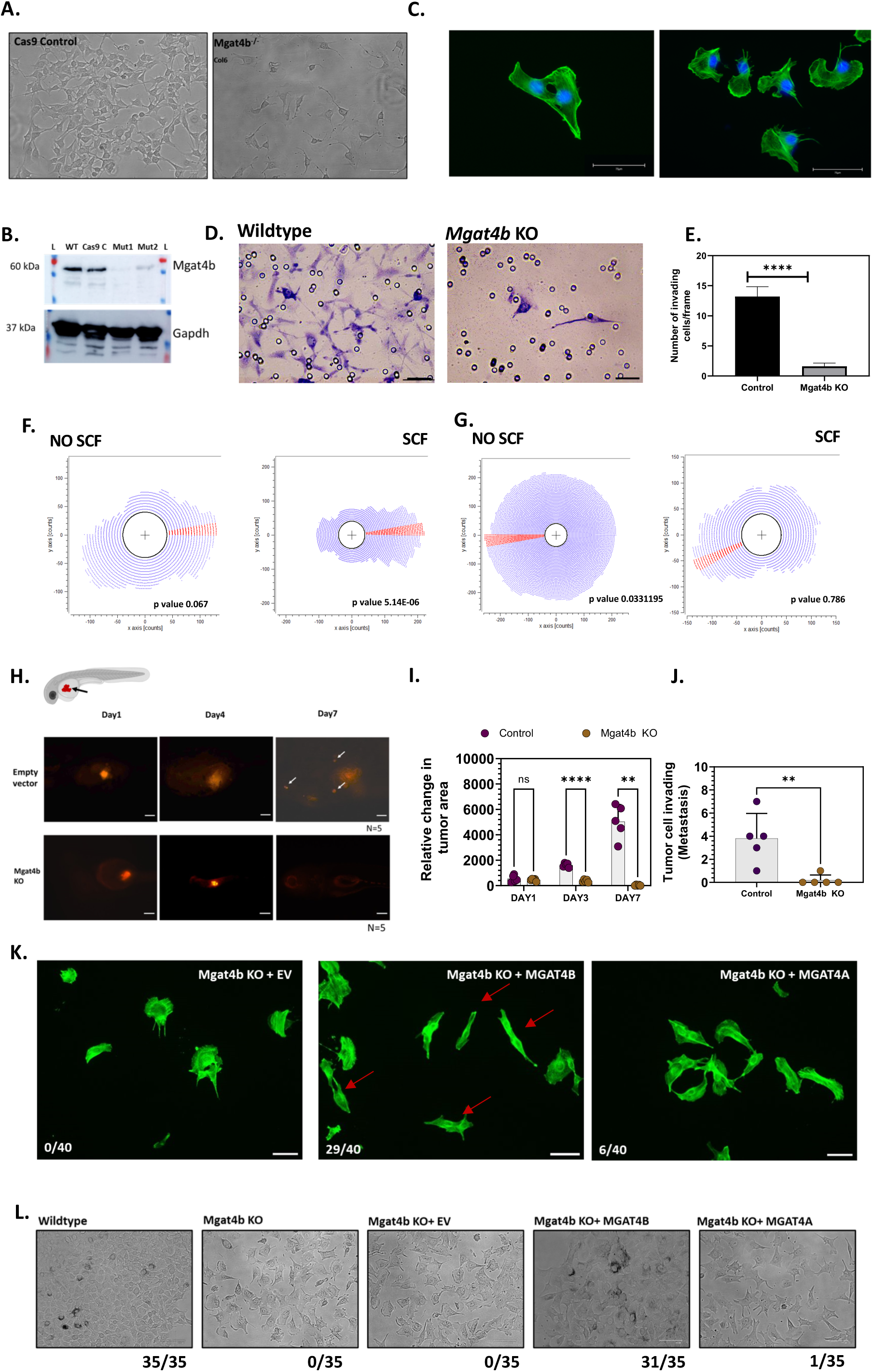
Loss of *Mgat4b* leads to reduced invasion capability and affects directionality of migrating melanocytes: **A.** Brightfield images shows wildtype control and *Mgat4b* knockout cells morphology, Scale bar-100 µm **B**. Western blot shows protein levels of *Mgat4b* in wildtype control and two clones of *Mgat4b* KO (Mut1 and Mut2) along with Cas9 control in B16-mouse melanoma cells **C.** Phalloidin staining shows actin filament distribution in wildtype control and *Mgat4*b knockout (Mut1) cells, Scale bar-75 µm **D.** Toluidine blue labels cells on Matrigel depicting the invasion capability of wildtype control and *Mgat4b* knockout(Mut1) cells, Scale bar-50 µm **E.** Bar graph depicting percentage of invasion of wildtype control and *Mgat4b* knockout cells (Mut1) **F, G.** Rose plot depicting spatial distribution of wildtype control and *Mgat4b* knockout cells(Mut1) in chemotaxis chamber assay **H.** Xenograft of mouse melanoma cells (wildtype control and *Mgat4b* knockout(Mut1) cells) into the 2dpf zebrafish yolk **I.** Bar plot shows relative change in tumor area of wildtype control and *Mgat4b* knockout(Mut1) cells, Scale bar-50 µm **J.** Bar plot shows number of tumor cells migrating out of primary cluster to invade neighboring tissues per frame **K.** Phalloidin staining shows actin filament distribution in *Mgat4b* KO(Mut1) cells complemented with Empty vector (EV), MGAT4B and MGAT4A protein, Scale bar-50µm **L.** Brightfield images showing cells morphology and colony formation in *Mgat4b* KO(Mut1) cells complemented with EV, MGAT4B and MGAT4A protein along with wildtype and *Mgat4b* KO cells (No transfection control), the numbers listed under each image indicate how many out of 35 cells exhibit the shown phenotype, Scale bar-175 µm *P ≤ 0.05, **P ≤ 0.01, ***P ≤ 0.001, ****P ≤ 0.0001 and ns P > 0.05

To evaluate if *Mgat4b* alters migration of mammalian melanocytes *in vivo*, we conducted xenograft experiments using wildtype and KO cells in zebrafish embryos. We studied the growth of DiI-labeled wildtype and KO cells by transplanting approximately 200-300 cells into the yolk sac of zebrafish embryos, 2 dpf. Images of the transplants were captured, and the cross-sectional areas were quantified at 4 and 7 dpf (Figure 4H). During the observation period, the cross-sectional areas of the KO xenografts gradually decreased, and by the end of 5 days, the KO cells had completely disappeared (Figure 4I). In contrast, control xenografts continued to grow and by day 5, the wildtype cells had begun to migrate and invade the surrounding tissues (Figure 4J). Hence, *Mgat4b* is essential for directionality and velocity of migration as well as growth of melanocytes *in vivo*.

### Profiling of differential glycosylation by lectin blotting

It is anticipated that these effects are a result of aberrant expression of glycoproteins, we suspected global alterations in *N*-glycosylation of cell surface proteins. To assess the proteins modified by *Mgat4b*, we employed *Datura Stramonium* Lectin (DSL), that selectively binds the branch generated by MGAT4 activity. We first performed lectin blot analysis on both wildtype and *Mgat4b* knockout B16 mouse melanoma cells, where the available cell quantity allowed for effective enrichment and subsequent target identification. We confirmed reduced DSL binding in the KO cell lysate compared to the wildtype, and this was supported by the FACS based assessment of cell surface labelling (Supplementary Figure 6E, F).

In the knockout cells we attempted to restore functionality by introducing exogenous MGAT4B, and included MGAT4A for comparison. Surface reactivity to DSL, an indicator of MGAT4B activity, showed that both isoenzymes restored levels comparable to wild-type cells. However, only MGAT4B was able to recover the filopodial phenotype (Supplementary Figure 6F, Figure 4K). Additionally, colony forming assays with rescued cells showed that MGAT4B, but not MGAT4A, restored the ’loose aggregate’ phenotype of MGAT4B knockout cells. Since only MGAT4B rescued both the filopodial and colony-forming phenotypes in knockout cells, this strongly indicates distinct substrate selectivity between these two glycosyltransferases (Figure 4K, L).

### Differential proteomics identifies target proteins of MGAT4B in melanocytes

Having established that *Mgat4b* is essential for melanocyte migration, we next sought to identify the cellular targets of glycosyltransferase responsible for this phenotype. We used a lectin-based approach to enrich glycoproteins specifically modified by MGAT4B in melanocytes. We assessed the levels of β1→4 linked GlcNAc sugars on cell surface of control and MGAT4B knockout zebrafish melanophores at 2 dpf using *Datura Stramonium* lectin (DSL) labelling. DSL has a strong affinity for β1→4 linked GlcNAc sugars, which are synthesized by MGAT4 activity (Varki et al., 2015). Flow cytometric analysis of surface accessible DSL reactivity was similar in other (non-GFP positive) cell types between the *mgat4b* mutant and NTC samples. However, in consistence with the earlier observed phenotype a two-fold reduction (p= 0.0271) was observed in GFP-expressing melanophores (Mitf+) in *mgat4b* mutant compared to control cells (Supplementary Figure 6 G-I). Melanocytes demonstrate high DSL reactivity, suggesting a requirement for MGAT4 activity in these cells. Interestingly, *mgat4b* mutant showed decreased melanocyte-specific DSL reactivity evidenced by the GFP and DSL double positive cells (Supplementary Figure 6E). Therefore, a DSL-based enrichment strategy would be an effective way to enrich for MGAT4B targets, allowing us to compare wildtype and *Mgat4b* KO cells to reveal the specific targets of this glycosyltransferase in melanocytes.

While DSL-based enrichment is an effective method for identifying MGAT4 targets in melanocytes, the low abundance of zebrafish cells presents a challenge. Therefore, for identifying MGAT4B targets, we opted to use B16 cells, where the knockout can be combined with DSL enrichment to analyse the glycoproteome affected by MGAT4B (Figure 5A). Differential proteomic analysis was carried out on DSL-enriched proteins from both wildtype and *Mgat4b* knockout B16 cells. Mass spectrometric analysis of DSL-enriched proteins consistently identified 104 proteins in wildtype and 58 proteins in knockout samples with at least one peptide being recognized across two biological replicate experiments. Among these 54 proteins were common between the two samples (Figure 5B, Supplementary table 2). After applying median filtering and removing common contaminant proteins and associated proteins such as ribosomal proteins, tubulins, and heat-shock proteins, we shortlisted proteins that are putatively modified by MGAT4B (Figure 5C). Based on literature analysis, these proteins may play a role in directional cell migration and invasion. Three of the shortlisted proteins, namely Glycoprotein non-metastatic melanoma protein B (GPNMB), Junctional Plakoglobin (JUP, also called as γ-catenin) and tyrosinase-related protein 1 (TYRP1) were further evaluated to confirm whether these proteins are substrates of MGAT4B that could explain the phenotypic outcomes seen in the knockout.

**Figure 5:**
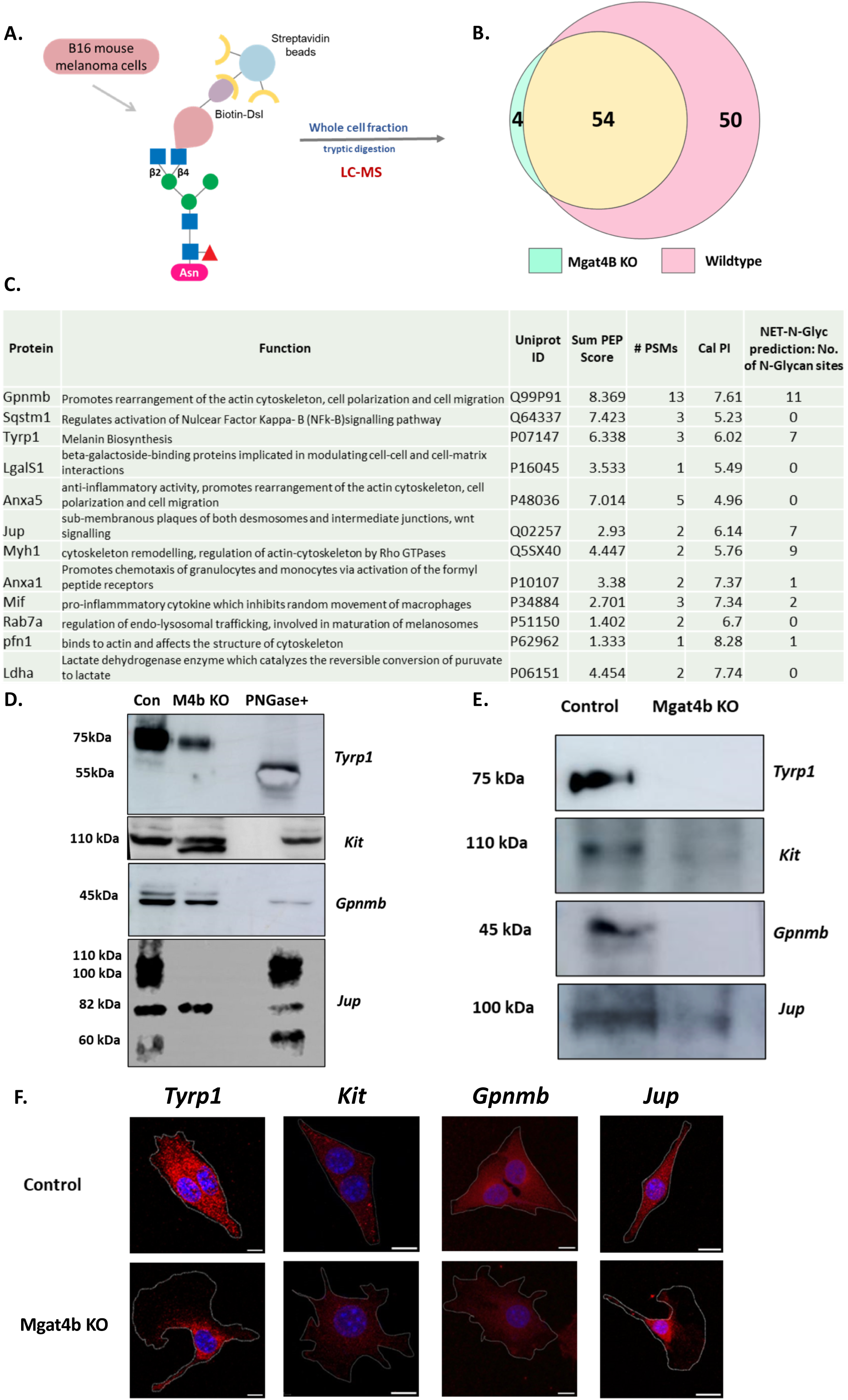
Differential proteomics reveal melanocyte specific target proteins of MGAT4B. **A.** Venn diagram depicting number of proteins identified by mass spectrometry analysis of the DSL-enriched fractions of Wildtype and *Mgat4b* KO proteins **C.** List of differentially enriched proteins in wildtype lysates with respect to *Mgat4b* KO along with their Uniprot ID, Sum PEP Score, PSMs, calc. PI and their functional profiling and predicted *N*-glycosylation sites by NetNGlyc -1.0 database **D.** Western blot analysis of *Tyrp1, Kit, Gpnmb* and *Jup* in wildtype and *Mgat4b* KO lysate along with PNGase treated wildtype lysate **E.** Western blot analysis of *Tyrp1, c-Kit, Gpnmb*, and *Jup* following biotinylated DSL pull-down in wildtype and *Mgat4b* knockout lysates **F.** Confocal image of immunocytochemistry using *Tyrp1, Kit, Gpnmb* and *Jup* (red) on permeabilized wildtype and *Mgat4b* KO cells, counterstained by DAPI(blue). The overlay of both channels is depicted. Scale bars: 50 µm

The transmembrane protein GPNMB is a known factor that promotes melanocyte growth and enhances extracellular matrix (ECM) binding (Tomihari et al., 2009). Its *N*-glycosylated ectodomain interacts with RGD-containing β1 integrin, leading to focal adhesion formation and subsequent activation of FAK and ERK pathways and controls rearrangement of actin-cytoskeleton, cell polarization and migration of melanocytes (Moussa et al., 2014; Tomihari et al., 2009). The protein Junction plakoglobin (Jup) is a member of the armadillo protein family, close homolog of β-catenin, and is a component of the desmosome connecting cadherins to cytoskeleton (Yin et al., 2005). Owing to the lack of signal peptide and its final destination in the cytoplasmic region, JUP does not seem to follow the secretory pathway. A NetNGlyc and NetOGlyc analysis of Jup protein sequence predicted seven and seventeen N-linked and O-linked glycosylation sites respectively, albeit no experimental data is available. TYRP1 is a melanosomal protein involved in the pigmentation process of melanocytes (Kobayashi et al., 1998). Our earlier data revealed that *Mgat4b* knockout cells were unable to respond to the SCF cue in their environment and failed to migrate towards it. Because of SCF’s critical role in guiding melanocyte movement, its receptor, Kit a known *N*-glycosylated protein essential for the survival of lateral and adult melanophores in zebrafish (Obata et al., 2014), was chosen for further validation. Therefore, in addition to the proteins identified in differential proteomics, we examined the tyrosine kinase receptor KIT as it was a compelling candidate for glycosylation modification by MGAT4B.

To determine whether these shortlisted proteins mediate the observed phenotypic effects of MGAT4B, we evaluated their protein levels, relative mobility on SDS-PAGE, ability to bind DSL, as well as the protein localization in wildtype versus *Mgat4b* KO cells. We performed western blots for each candidate with a reference of deglycosylated form using peptide-*N*-glycosidase F (PNGaseF). TYRP1 showed a mobility shift in tune with the known extensive *N*-glycosylation (NEGROIU et al., 1999). In *Mgat4b* KO cells mobility shift was not observed although the binding to DSL was completely abrogated (Figure 5D, E). However, a reduction in the levels of TYRP1 was observed (Figure 5D). This suggests that although MGAT4B-mediated modification does not significantly affect the overall glycan load, it remains essential for protein stability. Immunolocalization studies further confirmed a decrease in the overall protein levels (Figure 5F). In the *Mgat4b* knockout lysate, the Kit receptor showed two distinct bands around 110 kDa, whereas the wildtype and PNGase-treated lysates displayed only one band of the same size corresponding to the top band observed in the knockout. This suggests a possible proteolytic cleavage of the Kit receptor occurs in *Mgat4b* knockout cells, possibly impairing its ability to respond to the ligand SCF. Immunofluorescence suggested a decrease in the overall KIT levels in the KO and also its localization was non-uniform and restricted to regions around the nucleus. Surface accessibility could not be checked as FACS antibodies recognizing extracellular epitope of KIT did not show immunoreactivity in B16 cells. GPNMB antibody recognised two bands migrating closely around 45 kDa, with the upper band being diminished in the *Mgat4b* knockout lysate and completely disappearing in the PNGase-treated sample. DSL lectin recognized the upper band, indicating GPNMB to be modified by MGAT4B, and this modification is essential for stable expression of this glycoform (Fig 5D &E). Immunofluorescence also indicated an overall decrease in GPNMB expression in the knockout cells (Figure 5F). JUP consistently exhibited three additional bands 110, 100 and 60 kDa along with the main 82 kDa band (Figure 5D). DSL recognizes 100kDa form which is missing in *Mgat4b* knockout (Figure 5E). The striking absence of 100 kDa band for JUP in *Mgat4b* knockout suggests that JUP has a high probability of non-redundant dependency on MGAT4B for elaboration of the predicated seven *N*-glycan sites. The selective identification of glycoproteins, such as GPNMB, KIT, and JUP which are part of desmosome and adherens junctions, emphasize the role of protein-glycan interactions apart from the well-studied protein-protein interactions in cell-cell interactions.

### MGAT4B is a critical factor in melanoma progression

Having demonstrated the pivotal role of *mgat4b* in early melanocyte development, we set out to assess its potential implications in melanoma tumor initiation. Melanoma cells exhibit a dedifferentiated state similar to stem cells, where they possess heightened plasticity and tumorigenic potential (Brombin & Patton, 2024). Given Mgat4b’s involvement in orchestrating crucial developmental processes, including cell migration and patterning, there arises a compelling rationale to explore its function within the context of melanoma. Understanding the role of Mgat4b in melanoma may offer valuable insights into the mechanisms underlying tumor progression, metastasis, and perhaps even provide novel avenues for therapeutic intervention aimed at targeting the dysregulated glycosylation pathways characteristic of cancer cells.

To investigate the involvement of Mgat4b in melanoma, we initially examined its expression in skin cutaneous melanoma (SKCM) using the cancer genome atlas (TCGA) database. Remarkably, we observed elevated levels of MGAT4B in melanoma patients compared to normal individuals (Figure 6A). Subsequently, by analyzing two distinct datasets that categorized melanoma patients into metastatic and primary stages, we observed a significant increase in MGAT4B expression in metastatic patients compared to those at the primary stage (Supplementary Figure 7A), suggesting that MGAT4B may mediate aggressive disease in melanoma. Further exploration into the genetic alterations affecting MGAT4B revealed that ∼10% of melanoma patients harbour amplifications in MGAT4B, which could explain aberrantly high levels of MGAT4B expression in affected individuals (Supplementary Figure 7B). Molecular function enrichment analysis of melanoma, stratified by MGAT4B levels, revealed that among other pathways growth factor binding, cytoskeletal protein binding and integrin receptor binding were enriched (Supplementary Figure 7C). Importantly, patients with high MGAT4B expression have poor survival rates, exceeding those of patients with the common BRAF^V600E^ mutation. Our investigation further revealed that melanoma patients with both the BRAF^V600E^ mutation and elevated MGAT4B levels have significantly worse survival outcomes compared to those with only the BRAF^V600E^ mutation (Supplementary Figure 7D). These findings show the association between heightened MGAT4B levels and the aggressive nature of melanoma, emphasizing its significance as a prognostic indicator for patient outcomes.

**Figure 6:**
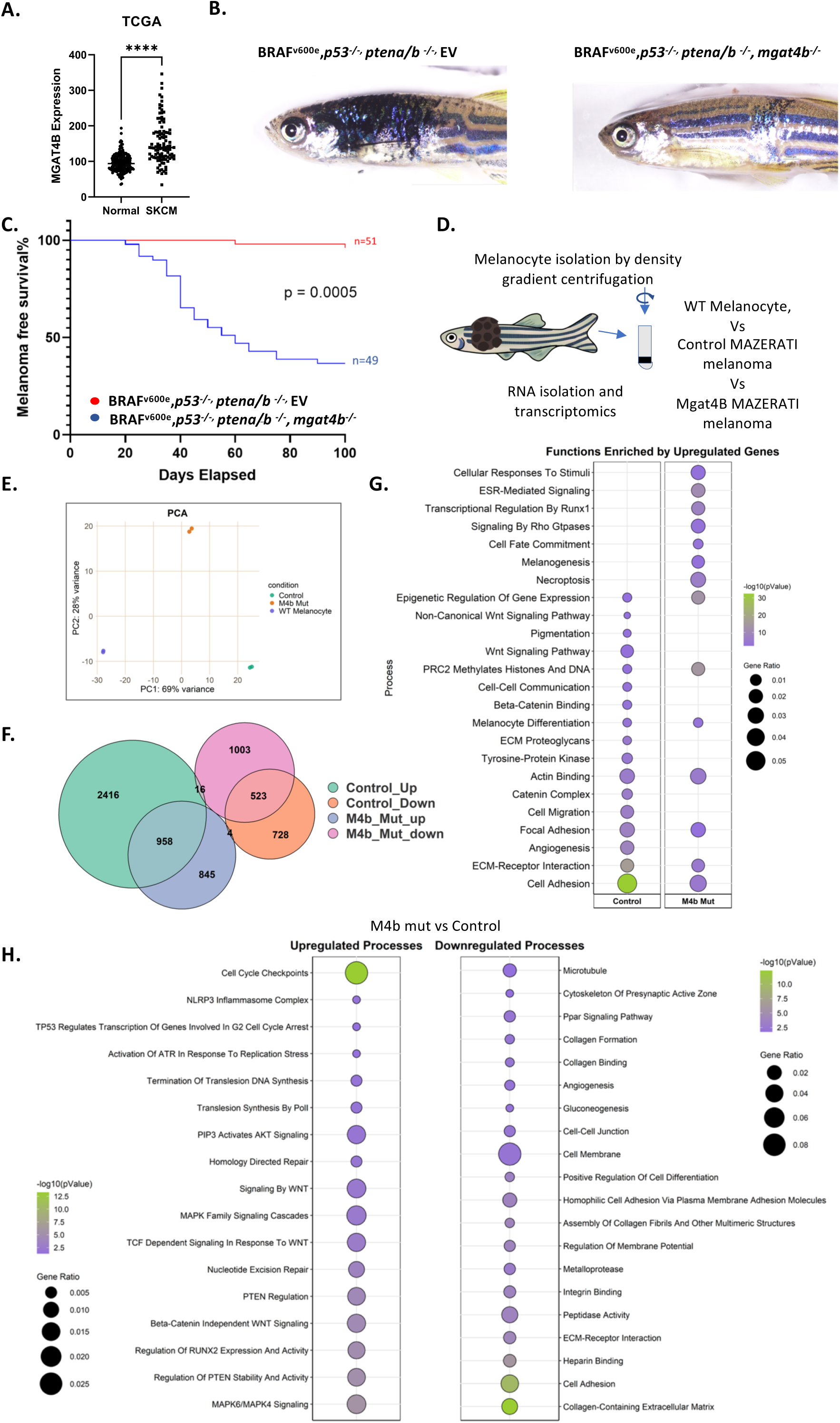
Elevated MGAT4B levels correlate with poor patient survival and are crucial for initiating primary tumors. **A.** Scatter plot depicting expression of MGAT4B in Skin cutaneous melanoma (SKCM) patients with respect to normal individuals, data procured from TCGA **B.** 60 days old Mazerati zebrafish displaying melanoblast-derived tumor developed by combinations of vectors expressing the oncogene BRAF^V600E^, targeting *tp53* and *ptena/b* initiate melanoma left-Mitfa-BRAF^V600E^, Mitfa-*p53*^-/-^, Mitfa-*ptena/b*^-/-^ and Mitfa:EV, right-Mitfa:*mgat4b*^-/-^ **C.** Melanoma-free survival curves of ASWT zebrafish injected with the indicated combinations of vectors expressing Mitfa-BRAF^V600E^, Mitfa-*p53*^-/-^, Mitfa-*ptena/b*^-/-^ and Mitfa:EV or Mitfa:*mgat4b*^-/-^ **D.** Schematic illustration of the experimental approach adapted to isolate and enrich(percol based gradient) melanophores from one month old wildtype and Mazerati fishes **E.** PCA analysis plot depicting relationship between the three samples sequenced **F.** Euler plot depicting the number of differentially expressed genes in melanocytes/melanoma cells from Mazerati fishes (control and M4b mut melanoma) with respect to wildtype melanophores **G.** Dot plot depicting functional enrichment from the upregulated genes in control and M4b mut melanophores/melanoma cells with respect to wildtype melanophores **H.** Dot plot depicting upregulated and downregulated processes in M4b mut melanoma with respect to control melanophores/melanoma *P ≤ 0.05, **P ≤ 0.01, ***P ≤ 0.001, ****P ≤ 0.0001 and ns P > 0.05

We utilized the well-established Mazerati model to genetically induce melanoma tumors in zebrafish, both with and without targetting *mgat4b* (Ablain et al., 2018). This approach, leveraging the MiniCoopR vector, allowed us to precisely express oncogenes and deactivate candidate tumor suppressor genes within zebrafish melanocytes. As previously established, we injected a cocktail of four plasmids to induce melanoma. This included plasmid expressing the BRAF^V600E^ oncogene and sgRNA targeting the tumor suppressors *p53* and *ptena/b*. We used Mitfa:Cas9;Mitfa:gfp plasmid to achieve melanocyte-specific gene ablation and melanoma initiation. Remarkably, the control fishes expressing non-targeting sgRNA rapidly developed aggressive tumors within a month, whereas the *mgat4b* mutant animals failed to develop melanoma altogether. Although melanocytes in the *mgat4b* mutant group showed signs of aggregation, they failed to initiate tumor formation demonstrating the indispensable role of *mgat4b* in initiating melanoma tumors in this zebrafish model (Figure 6B, C).

We explored the mechanisms that prevent mutant cells from initiating melanoma. To do this, we isolated mature melanophores from wild-type zebrafish and melanophores/melanoma cells from the skin of one month old MAZERATI zebrafish with and without *mgat4b* mutations. Cell isolation was done using density gradient centrifugation, followed by RNA extraction and sequencing (Figure 6D). In the wild-type group, we, enriched for mature, non-transformed melanocytes, while the control and *mgat4b* mutant group included both transformed melanoma cells and melanocytes. Principal Component Analysis (PCA) of the sequencing data showed clear segregation between the three groups, confirming that the *mgat4b* mutant (M4b mut) cells are distinct from both untransformed wild-type melanocytes and control melanoma cells (Figure 6E). While there was some overlap in gene expression between control and M4b mutant melanoma cells, most differences were unique (Figure 6F). In control melanoma cells, cell adhesion and migration processes were significantly enriched, but these were not similarly enriched in M4b mutant melanoma cells (Figure 6G). Consistent with scRNA seq and other experimental data discussed above, compared to control melanoma, M4b mutants showed upregulation of cell-cycle checkpoint genes and downregulation of cell adhesion and collagen-related extracellular matrix components (Figure 6H, Supplementary Figure 8A).

To validate this observation, we resorted to the B16 *Mgat4b* KO cells. We performed the complete proteomic analysis of B16 WT and B16 *Mgat4b* KO cells. Differential analysis of this proteome showed a similar enrichment of cell junction assembly, focal adhesion, and cell migration pathways (Supplementary Fig 8B, Supplementary Table 3). The activation of cell-cycle checkpoint genes may additionally contribute to the reduced melanoma growth in the zebrafish mutant. Reinforcing the role of MGAT4B in regulating these processes in melanocytes across model systems.

These observations established that ablation of complex glycosylation, such as that mediated by *mgat4b*, could be a viable strategy to deter the initiation of melanoma. To address whether targeting this complex *N*-glycosylation deters the already initiated melanoma we treated the MAZERATI animals with kifunensine, an inhibitor of α-mannosidase I, an enzyme crucial for trimming mannose residues during *N*-glycan processing (Elbein et al., 1991). By blocking this enzyme, kifunensine prevents glycoprotein maturation, causing the accumulation of high-mannose type *N*-glycans (Elbein et al., 1991). Since the MGAT4 enzyme requires a properly trimmed glycan substrate to add *N*-acetylglucosamine (GlcNAc) during the complex glycan processing pathway, kifunensine indirectly reduces MGAT4 activity by limiting the availability of suitable glycan substrates for its function. After melanoma initiation, once aggregation became visible, we administered kifunensine (150 µM). This treatment arrested melanoma growth, as indicated by the reduction in percent increase in melanoma area, suggesting that complex *N*-glycosylation remains an actionable target even after the initiation of melanoma (Supplementary Figure 9 A,B). To selectively inhibit MGAT4B without impacting other MGAT enzymes, we identified FDA-approved drugs with a higher binding affinity for MGAT4B. A virtual screening highlighted several compounds with a significant binding advantage (binding affinity difference > -1 kcal/mol) for MGAT4B compared to related MGATs (Supplementary Figure 9 C-E). In future, this approach could yield novel therapeutic intervention selectively targeting MGAT4B for use in deterring melanoma growth.

## Discussion

The role of *N*-glycosylation in melanocyte development is understudied, and although explored in melanoma, the causal relationship has remained unclear (Herraiz et al., 2011; Petrescu et al., 2003; Roux & Lloyd, 1986; Valencia & Hearing, 2012). We discovered that melanocyte precursors of NCCs lacking *mgat4b* activity show impaired migration, disrupting the formation of McSc pool, the subsequent source of melanocytes. Hence this selective loss leads to a decrease in the number of lateral melanophores and subsequently reduces overall melanocyte survival during metamorphosis (Mort et al., 2015). The reduction in melanocyte precursors is confined to a specific sub-population of migratory cells characterized by the expression of several lectins and cell adhesion proteins. It is likely that these cell surface *N*-glycoproteins could intricately guide the migrating cells effectively to their destination by specific interactions with the changing milieu of extracellular matrix enroute and facilitate establishment of the McSC pool.

In the context of melanoma, higher MGAT4B expression correlates with lower overall survival in patients with the most common driver mutation BRAF^V600E^, pointing to a probable role of elevated MGAT4B levels in promoting tumor aggressiveness (Castellani et al., 2023). When we attempted to initiate primary tumors in zebrafish by expressing BRAF^V600E^, the melanoma tumors did not develop in the *mgat4b* mutant, confirming its essential role. Along with the death of migrating melanocyte precursors and the resulting reduction in McSCs during development, MGAT4B alteration affects transformed melanocyte ability for cell adhesion and interaction with the matrix. While the cell adhesion changes could be attributed to the glycosylation of GPNMB and expression and localization of JUP, it is further exacerbated by changes in several cell adhesion genes as detected by RNA sequencing. This likely hinders the initiation of melanoma tumors, as the transformed cells form loose aggregate and fail to develop into melanoma tumor effectively. The other prominent effect of *mgat4b* mutation is the induction of cell cycle checkpoint that eventually deters the aggregated cells from further proliferation. Strikingly, same processes controlled by Mgat4b during melanocyte development also turn up during melanoma initiation reinforcing the recapitulation of developmental events in melanomagenesis.

Among the validated *N*-glycoprotein substrates of MGAT4B, GPNMB is the most promising candidate as this protein has been implicated in melanocyte development and also in melanoma (Han et al., 2021; Taya & Hammes, 2018; Tomihari et al., 2009). A substantial pool of GPNMB localizes to melanosomes, while a smaller but notable amount is found on the surface of melanocytes where it binds to Integrin beta1 *via* the RGD motif (Tomihari et al., 2009). Gpnmb is a bonafide *N*-glycoprotein that is a strong candidate substrate for MGAT4B, and is a MITF-dependent gene that has been identified as an effective target for preventing recurrent melanoma in patients treated with BRAF and/or MEK inhibitors (Han et al., 2021), (Rose et al., 2016). The phenotypic effects observed upon MGAT4B targeting can be explained by the alterations in GPNMB. Additionally, JUP is a likely *N*-glycosylated protein target of MGAT4B. Alternately as a component of desmosomes, JUP could link *N*-glycosylated MGAT4B target transmembrane cadherins desmogleins and desmocollins to intermediate filaments of the actin cytoskeleton (Kowalczyk & Green, 2013).

*N*-glycosylation is considered a ubiquitous process across all cell types, with selectivity typically being less emphasized. MGAT4B is under the regulation of MITF, is strikingly similar to the transcriptional control of the other isozyme MGAT4A by Foxa2 and Hnf1a in pancreatic beta cells (Ohtsubo et al., 2011). These two isozymes are non-redundant in the melanocyte phenotype, as their protein targets are very likely distinct. This immediately suggests a mechanism for cell-type specific regulatory events in glycosylation. The regulatory mechanisms as well as the downstream targets of these selective glycosyltransferases are distinct rendering MGAT4B to be a good therapeutic target for melanoma. In a previous systematic study DSL reactivity was found to be among the top differential glycome between primary and metastatic melanoma (Agrawal et al., 2017). The authors focused on FUT8 and its glycoprotein targets, including L1CAM, that were shown to play a crucial role in melanoma metastasis by affecting cell migration and invasion, making them potential therapeutic targets. Our complementary strategy attributes a causal role of *mgat4b* mediated glycosylation in the initiation of melanoma. As a proof of principle we demonstrate using Kifunensine, an inhibitor of complex glycosylation to be effective in reducing melanoma in the zebrafish model. Hence, targeting complex *N*-glycosylation branching, which is crucial for cancer initiation, presents a promising approach for melanoma treatment. While kifunensine demonstrates efficacy in melanoma reduction, its broader impact on *N*-glycosylation also results in higher toxicity (Elbein et al., 1990). We propose that a drug specifically targeting MGAT4B activity could be more effective in combating melanoma.

## Materials and Methods

### Zebrafish strains and maintenance

A zebrafish (Danio rerio) breeding colony was maintained at 28.5°C under a 14-hour light/10-hour dark cycle. The wild-type strain Assam WT (ASWT) and Tyrp1 reporter, Tg(ftyrp1: GFP) zebrafish lines were used for morpholino injections and were maintained according to standard zebrafish husbandry protocols. ftyrp1: GFP plasmid was a kind gift from Dr Xiangyun Wei (Zou et al., 2006), University of Pittsburgh School of Medicine, and the transgenic line as reported previously (Raja, Gotherwal, et al., 2020)was created at CSIR-IGIB zebrafish facility using tol2 transposase microinjections in ASWT line. Phenylthiourea (PTU;0.003%) was added to embryo water before 24 hpf to prevent melanin from masking the GFP fluorescence.

### Imaging

Bright-field imaging for live imaging and gene expression pattern (WISH) studies of zebrafish embryos was performed using a Zeiss stereo microscope (Stemi 2000-C). A Zeiss Axioscope A1 Microscope (with AxiocamHRc) was used for fluorescence imaging. Zeiss proprietary software was used to capture images, which were processed and analyzed in ImageJ software. Epinephrine treatment was given to adult fishes as described previously (Usui et al., 2018) and then imaged.

### Morpholino knockdown of zebrafish *mgat4b*

Antisense morpholino was synthesized from Gene Tools against *mgat4b* of zebrafish. The splice-block morpholino used in the study is ACATTTTGATCTTACCTCCGCAATG. Standard control was obtained from Gene-tools and injected at same dosage as *mgat4b* morphlino.

### Estimation of melanin content

Dorsal images of Control and *mgat4b* morphant embryos were taken at 2 dpf. The images were imported to ImageJ, and mean gray values were taken for melanophores of each embryo set. Mean gray values are inversely proportional to the melanin content of the cell. Corresponding values were then plotted using GraphPad Prism.

### *mgat4b* CRISPR mutant generation in zebrafish

The design of sgRNAs targeting *mgat4b* was conducted utilizing the CRISPRscan online tool. Candidates with a score exceeding 50 in CRISPRscan were selected. Subsequently, these chosen sgRNAs were assessed for potential off-target effects in zebrafish using the CasOFFinder online tool. Two or three of the highest-scoring CRISPRscan candidates, which exhibited no off-targets with ≤ 3 mismatches, were ultimately chosen for further investigation.

Chosen sgRNAs were annealed with the universal reverse primer provided by the CRISPRscan tool and then amplified. PCR purification of the sgRNA amplified product was performed using Qiagen PCR purification kit. IVT (in vitro transcription) of the PCR purified product was performed using Invitrogen T7 Megascript kit. The sgRNA+Cas9 cocktail was microinjected in zebrafish 1 cell stage embryos. The cocktail used for standard injections was: sgRNA – 55pg/nL, Cas9 – 330pg/nL, KCl – 330mM. These injected embryos were tested for the efficiency and presence of mosaic mutations using high-resolution melt analysis (HRMA) assay. For HRMA assay, from each injection batch, DNA was isolated from 8 random embryos and used as a template to set up a PCR with primers flanking the PAM site of each sgRNA. The presence of mutation will lead to formation of heteroduplexes during the PCR which will have low melting temperature compared to the non-injected control (NIC) embryos. Only those batches (putative Fos) were grown for each which had the presence of mosaic embryos in at least 2 out of 8 randomly selected embryos. The putative Fos were grown for each selected sgRNA. After these embryos attained the breeding age, they were set up for breeding with wildtype embryos. For each cross, the offspring embryos were collecting and at 5 days post fertilisation, DNA was isolated from randomly 8 embryos and high-resolution melt analysis (HRMA) was performed to confirm the F0s. The clutch in which there was observable HRMA peak shift or a CT difference of ≥ 0.5 compared to the wild type embryos were set up for growing as putative F1s. After these putative F1 fishes were grown and reached breeding stage (3 months old), they were crossed with wildtype fish and their fins were clipped after breeding. DNA isolated from the fins was used as a template for HRMA to confirm the presence of heterozygosity in the embryos and confirmation of the F1 parent fish genotype. The crosses for which HRMA came out to be positive were grown as the F2 batches to obtain homozygous mutants.

### Melanophore-specific CRISPR mutant generation

The cell-type specific knockout strategy was adapted using MinicoopR vector in which Mitfa promotor (melanocyte specific promotor) drives Cas9 and mitfa mini gene and has space for two sgRNA under Ubiquitous (U6) promotor (Ablain et al., 2018). Mitfa mini gene was replaced with GFP using restriction digestion approach. Mitfa: Cas9, Mitfa: gfp plasmid was injected a little ahead of one cell stage in zebrafish embryo and imaged at 2dpf stage. Two sgRNAs were designed using CHOPCHOP (https://chopchop.cbu.uib.no/) targeting *mgat4b* gene. The cloning was performed by digesting the plasmid using BseRI enzyme. We confirmed the clones using PCR based approach using Forward primer specific to plasmid and sgRNA as a reverse primer. The PCR validated clones were sanger sequenced and then injected at one-cell stage in Zebrafish embryos and the imaging was done at various developmental stages.

### Flow cytometry

Cell counts for Mitfa:gfp cells and ASWT embryos (for melanophore counts) were performed using an BD FACSAria II instrument flow cytometer. Briefly, embryos were dechorinated using pronase (5 mg/ml) (Sigma-Aldrich, P8811) for 10-15 min and collected in microfuge tubes. The embryos were deyolked in ice cold Ringer’s solution using a micropipette tip and spun at 100 g for 2 min in a tabletop centrifuge at 4°C (Eppendorf, 5418R). The supernatant was discarded and the embryo bodies were trypsinized using TrypLE Express (Thermo Scientific, 12604039) for 15 or 30 min for 24 hpf or 48 hpf, embryos respectively, at room temperature. The cell suspension was passed through a 70 μm cell strainer and washed twice with ice cold phosphate-buffered saline. The cell suspension was analyzed and sorted using the imaging flow cytometry system.

#### Surface labelling of zebrafish cells

The cell suspension was incubated with the APC or FITC labelled Datura Stramonium lectin (BioPLUS GlycoMatrix DSL-Texas Red (21761014-1), DSL-Fluorescein (FL-1181-2)) for 1 hour on ice and washed three time for 5 minutes each. Cells were analysed using BD FACSAria II instrument flow cytometer and later analysed using FlowJo^TM^ (TreeStar).

### B16 mouse melanoma culture

The B16 mouse melanoma cell line was cultured in DMEM-high glucose medium (Gibco, Life Technologies) supplemented with 10% fetal bovine serum (FBS; Gibco, Life Technologies). The cells were maintained in a 37°C incubator with 5% CO₂.

### Phalloidin staining

Cells cultured to a confluence of 50,000 cells per 6 well plate for 20 mins at 37°C. Immediately after the incubation the cells were washed with 1x DPBS, twice and then fixed with 4% paraformaldehyde at 37°C. Once fixed the cell were either stored in DPBS at 4°C or taken ahead for F-actin staining with Phalloidin (Alexa Fluor 568 Phalloidin-A22283, ThermoFisher scientific). Phalloidin stained coverslips were then imaged at 20X or 40X with EVOS M7000 Imaging System, Invitrogen.

### Colony formation assay

For the colony formation assay, B16 cells (control and Mgat4b ko cells) were seeded at a very low density of 100 cells/cm2 and cultured for appropriate number of days as per the experimental requirement, for a maximum of 7 days. Colonies formed from a single cell were imaged at day7.

### Immunofluorescence

The cells plated on coverslips were washed twice with 1× PBS and then fixed with 4% PFA at 37°C for 20 min. The cells were again washed twice with 1× PBS and permeabilized with 0.01% Triton X-100 (Sigma). The cells were blocked with 5% normal goat serum (Jackson Laboratories) overnight at 4°C. The cells were washed twice with PBST (1× PBS with 0.01% Tween-20). The cells were then incubated with 1:100-250 dilution of Tyrp1 (abcam, ab178676), Kit (thermo, 14-1172-82), Gpnmb (abclonal, A1427), Jup (Gamma catenin, PG-11E4) in a moist chamber for 1 h at room temperature. Incubation with secondary antibody Alexa fluor 594 Molecular probes, Thermoscientific) was performed at room temperature for 1 h. The cells were again washed with PBST. The cells were then mounted on slides using Antifade slow-fade DAPI (Molecular probes, Invitrogen) and visualized using Confocal Microscope (Leica SP8).

### Western blotting

Cells were trypsinized with 0.1% trypsin and the pellet was washed twice with 1× phosphate-buffered saline (PBS; Gibco Life Technologies). NP-40 lysis buffer (Invitrogen) was added to the pellet and was incubated on ice for 30 min with pipetting at interval of 10 min. The cells were centrifuged down at 11,200 g for 30 min at 4°C (Eppendorf Centrifuge 5415 R). The supernatant was collected and transferred to a fresh microfuge tube. The protein was estimated using standard BCA protocol (Pierce BCA protein assay kit; Thermoscientific). Equal amount of protein from each sample were resolved in 10 or 12% % SDS gel in 1× Tris-glycine buffer. The gel was blotted onto 0.45 µm PVDF membrane (Millipore) at 150 mA for 1 h. 5% Skim milk was used for blocking for 1 h at room temperature. Incubation with primary antibody was performed for overnight at 4°C. Primary anti-body Tyrp1 (abcam, ab178676), Kit (thermo, 14-1172-82), Gpnmb (abclonal, A1427), Jup (Gamma catenin, PG-11E4), Mgat4b (abclonal, A12810). After washing the blot with 1× TBST, the blot was incubated with HRP-conjugated secondary antibody for 1 h at room temperature. After washing with 1× TBST, the blot was developed using ImageQuant™ LAS 500 chemiluminescence instrument. Densitometry analysis was performed using ImageJ software.

Zebrafish lysates were as previously described (Liu et al., 2009)and SDS-PAGE was performed as described above.

### Reanalysis of 5dpf Sox10+ scRNA sequencing data

Single cell RNA sequencing data of 5dpf sox10+ zebrafish cells capturing multiple sox10 expressing neural crest lineages, including melanophores. Data was analyzed using Seurat v4 and mimics the authors analysis. (**Ref: GSE131136)**

### Single-cell sequencing and analysis

Mitfa+ve cells were sorted by FACS from 36 hpf zebrafish. The single-cell RNA libraries were prepared for both samples using the 10× genomics Chromium Next GEM Single Cell 30 Reagent Kit v3.1. The library QC and quantification was done using Agilent bioanalyzer HS DNA kit. The libraries were pooled and sequenced on the NextSeq 2000 platform. Raw bcl files were converted to final count matrix using Cell Ranger v6.1.2 software following the tutorial provided on the 10x genomics website.

FastQ files from control and *mgat4b* knockout zebrafish Mitfa+ve cells were aligned using the 10x Genomics CellRanger v7.2 (Zheng et al., 2017) pipeline to zebrafish genome (Ensembl GRCz11). The Gene-cell matrices (Control: 1923, *mgat4b* knockout: 4120) were uploaded on RStudio (R version 4.3)(Ihaka & Gentleman, 1996) and standard quality control metrics with the Seurat package (v.5.1) (Satija et al., 2015). Only cells with total features >200 and mitochondrial gene counts (%) < 5 were considered as high quality and kept for further analyses. Random sampling was performed in *mgat4b* knockout sample to have the same number of cells as in Control sample for comparison between the two samples. The data from both the sample were integrated using 2000 integration anchors with the help of Find Integration Anchors and Integrate Data functions to merge the datasets and remove any possible batch effects. Then, the Louvain clustering of the integrated dataset was performed with Seurat (v.5.1) using the FindNeighbors and FindClusters functions (dims = 13, resolution = 0.07) after performing linear dimensionality reduction and checking the dimensionalities of the datasets visualized with elbow plots. Data were projected onto 2 dimensional spaces using Uniform Manifold Approximation and Projection (UMAP) (Ghojogh et al., 2023) using the same dimensionality values listed above.

Cluster specific genes were identified using the FindAllMarkers and FindMarkers function in Seurat with default parameters (Wilcoxon Rank-Sum test that compares a single cluster against the others). Cluster annotations were given referring previously published datasets. Functional enrichment for markers from specific clusters were performed using GSEA in Clusterprofiler package(Yu et al., 2012) in RStudio.

Plots were generated either using Seurat or ggplot2 (Wickham, 2011).

Pseudotime trajectory construction was performed on the integrated dataset using Monocle3 (Cao et al., 2019; Qiu et al., 2017; Trapnell et al., 2014).

Cell-Cell communication analysis was performed using Nichenetr package (Browaeys et al., 2020). Briefly, the migratory MIX+ cluster was considered as recipient and receptors from this cluster was identified. All the remaining cells were considered as sender and ligands from these clusters were identified. As Nichenetr package is specific for human/mouse data, we used Diopt(Hu et al., 2011) tool to find orthologs of the receptors and ligands identified from our clusters. These orthologs were used to construct receptor-ligand interaction information for our dataset.

### Melanophore Enrichment from Zebrafish Skin and Melanoma Biopsies and RNA sequencing

Zebrafish skin and melanoma biopsies were collected and enzymatically dissociated using Liberase TM (Sigma Aldrich – 5401020001) at a concentration of 0.25 mg/mL in phosphate-buffered saline (PBS). The tissue was manually triturated to obtain a single-cell suspension, which was then filtered through a 70 µm cell strainer and centrifuged at 500 xg for 15 minutes at 4°C. The resulting pellet was resuspended in 2% Fetal Bovine Serum (FBS) in PBS and layered onto a 50% Percol solution (Higdon et al., 2013). Centrifugation was repeated at 500 xg for 15 minutes at 4°C. After careful removal of the supernatant, the enriched melanophores at the bottom were washed once with PBS, lysed with Trizol, and processed for RNA extraction.

### RNA Seq Library Preparation

RNA sequencing libraries were prepared using the Illumina TruSeq® Stranded Total RNA Library Prep Gold kit (cat. no 20020598) following the manufacturer’s protocol (1000000040499 v00). A total of 250 ng of RNA isolated from the enriched melanophores/melanoma cells from zebrafish skin was used as input. Cytoplasmic and mitochondrial rRNAs were depleted using the Illumina Ribo-Zero rRNA removal beads to reduce the abundance of these highly expressed RNA species.

Following RNA depletion, the RNA was fragmented and reverse transcribed to synthesize the first strand of cDNA. The RNA strand was then digested, and second strand cDNA synthesis was performed to create double-stranded cDNA. The 3’ ends of the double-stranded cDNA were blunted, and A-tailing was performed to facilitate ligation of index adapters. Subsequently, PCR-based amplification was performed to enrich the cDNA libraries.

Libraries were purified and size-selected using AMPure XP beads (Beckman Coulter, A63881). The quality of the libraries, including size distribution, was assessed using the Agilent HS D1000 Screen Tape (5067-5587) on an Agilent 2200 TapeStation, and library quantification was performed with the Qubit dsDNA High Sensitivity Assay Kit. Final libraries were diluted to 2 nM and pooled equimolarly. Sequencing was performed on the Illumina NextSeq 2000 platform using the XLEAP P4 300-cycle sequencing kit, generating paired-end reads (2 × 151 bp) with a final loading concentration of 650 pM.

#### Data analysis

Raw fastq files for each sample were first adaptor removed and trimmed for high confident base reads using Trimmomatic tool (Bolger et al., 2014). The trimmed files were further aligned with zebrafish reference genome GRCz11 using STAR aligner (v2.7.8)(Dobin et al., 2013). Featurecounts tool was used for calculation of raw counts for each gene in the sample (Liao et al., 2014). The differential gene expression analysis was performed using DESeq2 package (v1.40.2) in Rstudio(Love et al., 2014). Functional enrichment analysis was performed using DAVID webtool(Sherman et al., 2022).

### Live imaging of Zebrafish embryos

Live imaging of embryos between 28 hpf-36 hpf were done using Leica SP8 STED confocal microscope. Images were captured in xyzt mode at 10x magnification. The embryo was laid laterally to visualize dorso-lateral cell migration in trunk region. The sample preparation was done exactly as depicted by (Upadhyay & Papadakis, 2020).

### Annexin V and acridine orange assay

Zebrafish embryos were dechorionated using pronase (SIGMA; Roche). For deyolking, dechorionated embryos were collected in a 1.5ml microcentrifuge tube. 200µl of ice-cold Ringer’s solution was added to the embryos and mixed well by pipetting using a 200µl tip. The tubes were centrifuged at 100 rcf for 1 min, supernatant was removed and further 1ml of ice-cold ringer’s solution was added to the embryo body pellet. The suspension was mixed gently by inverting the tubes twice and centrifuged at 400 rcf for 1 min and the supernatant was discarded. A single cell suspension was prepared by adding 10ml TrypLE Express (ThermoFisher Scientific; 12604013) to the deyolked embryos in a fresh petri dish. The solution containing embryo bodies were mixed to decrease aggregation. The petri dishes containing the deyolked embryos were incubated at room temperature for 15 mins (<24 hpf) or 30 mins (24 – 30hpf) and were occasionally flushed with 1 ml pipette to aid disintegration of cells. A 70µm cell strainer was placed above a 50 ml falcon and the single cell suspension was passed through to remove cell clumps and other particles >70um. The cell suspension was flushed a few times with the same solution so as to remove the cells adhering to petri dish. The samples were centrifuged at 1500 rcf for 5 mins at 4°C in swinging bucket rotor mode. The supernatant was discarded and the pellet was resuspended in 1ml ice cold 1Xphosphate buffered saline (PBS). The cells were centrifuged again at 1500 rcf for 3-5 mins at 4°C; the supernatant was discarded and the pellet was resuspended again in 1X PBS. Harvested cells were treated with the suggested concentration of recombinant Annexin V Pro APC (EBioscience™, Cat No. BMS306APC-100) for 15 minutes, followed by washing with the binding buffer provided in the kit. Subsequently, the cells were analysed using FACS.

Zebrafish embryos were subjected to staining with the vital dye acridine orange to quantify the number of apoptotic cells per embryo. The assay involved immersing the embryos in a solution containing 10 µg/mL of AO (786-2074, G-bioscoences) in E3 media. After a staining duration of 60 minutes, the embryos underwent three consecutive washes in E3 media. Following staining, the embryos were transferred to Petri dishes for imaging purposes.

### Chromatin immunoprecipitation and qRT PCR

B16 Melanoma cells at 80% confluence was fixed with 10% formalin (Sigma Cat No HT501128) and incubated at 37°C for 10 min. 2.5 M Glycine was added to the cells and again incubated at 37°C for 10 min. Cells were washed with ice-cold 1× PBS containing protease inhibitors. Cells were then scraped and centrifuged at 112g for 5 min at 4°C. The cell pellet was lysed in SDS lysis buffer (1%SDS, 10 mM EDTA, 50 mM Tris (pH 8.1)) on ice for 30 min. The cells were then sonicated on bioruptor (DIAGENODE) in ice. The chromatin lysate was then estimated for protein content using BCA kit (Pierce). 10 µg of MITF C5 antibody (Abcam) was taken and incubated with Protein G or Protein A Dyna-beads (Thermo-scientific) overnight at 4°C on a rotator. The next day, the sera was cleared and washed with ice-cold dilution buffer (2 mM EDTA, 150 mM NaCl, 20 mM Tris HCl (pH 8)). 500 µg of chromatin lysate was added to the beads and the final volume was made up to 750 µl using the dilution buffer and incubated for 6 h at 4°C in a rotator.10% of the lysate was kept separately as input. After incubation, the magnetic beads were washed with low salt buffer (0.1% SDS, 1%Triton X-100, 2 mM EDTA, 20 mM Tris HCl (pH 8), 150 mM NaCl), high salt buffer (0.1% SDS, 1% Triton X-100, 2 mM EDTA, 20 mM Tris HCl (pH 8), 500 mM NaCl), and LiCl buffer (0.25 M LiCl, 1%Igepal C-630, 1 mM EDTA, 10 mM Tris HCl (pH 8), 1% deoxy-cholate). Finally, the magnetic beads were incubated overnight at 65°C in elution buffer (1% SDS, 0.75% sodium bicarbonate) and 1 µl of 20 mg/ml proteinase K (Sigma) for elution and subsequent reverse cross-linking. The magnetic beads were separated from supernatant and column purified using Qiagen PCR purification kit, the input control was also included in the purification step. SYBR qRT-PCR was setup using 5 µl of eluted DNA, and graphs were plotted fold enrichment.

### CRISPR based mutagenesis of Mgat4b in B16 mouse melanoma cells

Guide RNA for each target genes were designed with CRISPRscan online tool. Chosen sgRNAs were annealed with the universal reverse primer provided by the CRISPRscan tool and then amplified. PCR purification of the sgRNA amplified product was performed using Qiagen PCR purification kit. IVT (in vitro transcription) of the PCR purified product was performed using Invitrogen T7 Megascript kit.

The cocktail containing 625ng CRISPR-RNA, Cas9 nuclease-2500ng and lipofectamine-5ul was incubated at 25*c for 10min. RNP complex was added to the wells according to the labels. After 4-5 hours, the transfection was terminated by washing the cells with DPBS and add fresh complete media (DMEM High glucose with 10%FBS and 1x Anti-anti). The RNP transfected cells were harvested with trypsin-EDTA and resuspended in 1x PBS. Singlet Cells were then sorted into 96-well plates with the help of BD FACSAria™. Sorted cells that formed a visible single colony in each well within a week became ready to be passaged 2 to 3 weeks after sorting. These colonies were individually validated for the mutation using Sanger sequencing followed by western blotting.

### Modelling Approach in Zebrafish for Rapid Tumor Initiation (MAZERATI)

Primary melanoma was initiated in ASWT fishes using method discussed in (Ablain et al., 2018). Plasmids from Addgene 118846, 118850, 118841, 118845 were used for the study. These plasmids will overexpress BrafV600e oncogene specifically in melanocytes and target p53 and pten tumor suppressors again specifically in melanocytes. Addition to this we used modified Mitfa:cas9;Mitfa:gfp plasmid to target mgat4b in the melanocytes. Cocktail of five plasmids were injected in suggested concentration, in one-cell stage to transform the melanocytes. Aggressive tumors were initiated in the zebrafish within a month.

### Chemotaxis Assay

Chemotaxis chamber assay was set-up in according to the protocol described by the manufacturer. Briefly, each chemotaxis coverslip has 3 (1mm wide) trough regions where the cells are seeded. Each trough is surrounded by media reservoirs on either side. The trough is connected to media reservoirs on either side by a small opening that allows for gradual diffusion of chemotactic agent across the trough, generating a concentration gradient. Once the cells are adhered and the chemotactic agent is added into the respective reservoir, the chamber is set-up on the microscope stage for imaging. Growth promoting chemokine stem cell factor (SCF, PeproTech #300-07). Time-lapse XYT imaging was done for ∼20 hours with time interval of 30mins. During the imaging the slide was maintained at 37°C with 5% CO2 using Environment controlling unit. Imaging was done with Confocal (SP8) microscope at 10x magnification. Chemotaxis analysis was done by individually generating the tracks of each cell (>50 cell/per experiment) using manual tracking plugin in Fiji and then processing the track information in chemotaxis and migration tool (Fiji Plugin) developed by ibidi.

### Cell Invasion Assay

Cell invasion assay was performed using a matrigel assay system (Corning Biocoat) in a 6-well format. Briefly, 100,000 cells each for wildtype and mgat4b knockout were seeded onto Matrigel-coated invasion chambers as per the manual. The next day, cells were fixed and labelled by Toluidine Blue to stain the invaded cells. The invaded cells were imaged using a brightfield microscope with at least 4 to 6 fields per well, and ImageJ was used to quantify the number of invaded cells. Relative invasion was calculated by normalizing all data to the number of invaded cells in the control group.

### B16 mouse melanoma cells processing followed by LC-MS/MS

Cells were lysed using RIPA buffer and centrifuged at 15,000 g for 15 minutes at 4°C. The supernatant was transferred to a new microcentrifuge tube for proteomics analysis. Protein precipitation involved overnight incubation of the supernatant with four times volume of pre-chilled acetone, followed by centrifugation at 15,000 g for 15 minutes at 4°C. The resulting protein pellets were resuspended in 50 mM ammonium bicarbonate buffer (pH 7.8). Protein quantification was performed using the Bradford assay, and 20 μg of protein from each sample was used for quantitative proteomics analysis (SWATH-MS). In brief, protein samples were treated with PNGase F (V483A) and incubated at 37°C for 4 hours. Following this, samples were reduced by adding dithiothreitol (DTT) to a final concentration of 2 mM and heating at 56°C for 30 minutes. The samples were then cooled to room temperature and alkylated with iodoacetamide (IAA) at a final concentration of 2.2 mM, incubating in the dark for 15 minutes. The samples were digested with trypsin (V5111, Promega) at a 1:20 enzyme-to-substrate ratio for 18 hours at 37°C. Tryptic peptides were purified using Oasis HLB 1 cc Vac cartridges (Waters) according to the manufacturer’s protocol, vacuum dried, and stored at -20°C until LC-MS/MS analysis

#### Data acquisition

The tryptic digests from the samples were suspended in 0.1% formic acid and analyzed on a quadrupole-TOF hybrid mass spectrometer (TripleTOF 6600, SCIEX) coupled to an Eksigent NanoLC-425 system using the Sequential Window Acquisition of All Theoretical Mass Spectra (SWATH-MS) method by operating the mass spectrometer in data-independent acquisition mode. Optimized source parameters were used, curtain gas and nebulizer gas were maintained at 25 psi and 20 psi respectively, the ion spray voltage was set to 5.5 kV, and the temperature was set to 300°C. About 4 μg of peptides were loaded on a trap column (ChromXP C18CL 5μm 120 Å, Eksigent, SCIEX) and online desalting was performed with a flow rate of 10 μl per minute for10 min. Peptides were separated on a reverse-phase C18 analytical column (HSS T3, 100 Å, 1.8 μm, Waters) in 55 minute buffer gradient at a flow rate of 6 μl/minute using water with 0.1% formic acid (buffer A) and acetonitrile with 0.1% formic acid (buffer B) as follows:

| Time (min) | %A | %B |
| --- | --- | --- |
| 0 | 97 | 3 |
| 38 | 75 | 25 |
| 43 | 68 | 32 |
| 45 | 20 | 80 |
| 45.5 | 10 | 90 |
| 48 | 10 | 90 |
| 49 | 97 | 3 |
| 57 | 97 | 3 |

A SWATH-MS method was created with 96 precursor isolation windows, defined based on precursor m/z frequencies in DDA run using the SWATH Variable Window Calculator (SCIEX), with a minimum window of 5 m/z. Data were acquired using Analyst TF 1.7.1 Software (SCIEX). Accumulation time was set to 250 msec for the MS scan (400-1,250 m/z) and 25 msec for the MS/MS scans (100-1,500 m/z). Rolling collision energies were applied for each window based on the m/z range of each SWATH and a charge 2+ ion, with a collision energy spread of 5. Total cycle time was 2.79 s. **Data Analysis:** Raw file with .wiff extension from SWATH-MS runs were analysed using spectronaut software version 19 (Biognosys) using directDIA workflow. For protein identification, Mus Musculus protein database from uniprotKB with 54,707 entries was used with upto 2 missed cleavages. False discovery rate (FDR) was controlled at 1 % for proteins. Rest of the parameters were kept at Spectronaut Pulsar’s default settings, cross-run data normalization was performed where the normalization strategy was set to automatic. Quantitative data was exported in the form of ‘Run Pivot Report’ and differential protein analysis was performed in Microsoft Excel.

### DSL Blotting for glycan profiling

B16 cells (wildtype (WT) and *Mgat4b* KO) were washed with cold DPBS and trypsinized with 0.1% trypsin (Gibco, 15090046 diluted in Versene, Gibco, 15040066). and 10 million cells were counted for each sample. Cell pellets were resuspended in the RIPA buffer according to their size (G-Biosciences-786-490). Protein concentrations of total lysates of WT and *Mgat4b* KO cells were estimated by the Bradford assay (Da Silva & Arruda, 2006). Proteins were resolved through a 7.5 % SDS-PAGE and transferred to nitrocellulose membrane at a constant current of 250 mA for three hours at 4 °C. Membranes were blocked at room temperature with 2.0 % gelatin in 0.1 % PBST (phosphate buffered saline with 0.1 % v/v Tween-20) (10 mL, 1.0 h) followed by incubation with fluorescein isothiocyanate-conjugated Datura stramonium lectin (DSL) in PBST (1:5000 dilution; stock conc. 2.0 mg/mL; 5.0 mL, 1.0 h). Membranes were washed with 0.1 % PBST (10 mL × 6, five min incubation for each wash). The blots were visualized using a digital fluorescence imager (Amersham Typhoon) using a Cy2-filter for excitation. Two biological replicates, each with two technical replicates, were performed for the lectin blotting experiments.

### Proteomics study of DSL affinity enriched glycoproteins

Both B16F10 wild-type and KO cells were cultured in T-75 tissue-culture treated flasks under sterile conditions in DMEM supplemented with 10 % fetal bovine serum (FBS) and 1.0 % pen-strep (50 U/mL of penicillin and 0.05 mg/mL of streptomycin) in a humidified incubator with a 5.0 % carbon dioxide atmosphere at 37 °C. Once the cells attained 70-90 % confluency, the media containing the floating cells was aspirated. The adherent cells were gently washed with PBS (5.0 mL) once and aspirated. Cells were detached by incubating with 0.25% trypsin (1.0 mL) at 37 °C for 2-5 min and diluted with complete media (9.0 mL). Cells were collected in 15 mL tubes and centrifuged at 500g for five min, supernatants were aspirated, and discarded. Cells were re-suspended in PBS (1.0 mL), cell density was measured using a Z2 particle counter, cells were aliquoted in 1.5 mL microcentrifuges, centrifuged, and supernatants were aspirated. Both wide-type and KO B16F10-cells (5.0 × 106 cells) were suspended on lysis buffer (10 mM tris base, 140 mM NaCl, 1.0 % v/v triton-X-100, and 1.0 mM phenylmethylsulfonyl fluoride in MS-grade water, pH 7.4) (100 mL), probe sonicated (amplitude 25 %; Sonics, Vibra-cell) for 15 min on an ice-bath (amplitude 25 %), centrifuged (16000×g, 10 min, 4 °C), and the supernatant were transferred to fresh pre-labelled tubes. Protein concentrations were estimated using the Bradford assay. Total cell lysates (1.0 mg in 500 mL PBS) were allowed to bind to biotin-conjugated DSL (b-DSL) (5.0 mL from a stock solution containing 2.0 mg/mL) overnight at 4 °C with gentle agitation on an end-to-end rotor. The mixtures were treated with streptavidin-conjugated dynabeads (50 mL of slurry; binding capacity of 10 mg biotinylated antibody/mg beads; pre-washed with PBS (500 mL, thrice)) and incubated at 4 °C for two hours with gentle mixing. Samples were placed on the magnetic separator for 1.0 min, the supernatants were aspirated and discarded. The beads were washed in NP-40 buffer (0.1 % v/v NP-40 in PBS; 100 mL; three times) and the supernatants were aspirated and discarded. The beads were re-suspended in 200 mM N-acetyl-D-glucosamine (GlcNAc) solution in PBS (50 mL), vortex mixed for 10 min at room temperature, placed on the magnetic separator for 1.0 min, and the supernatant was carefully aspirated. Beads were subject to a second round of elution with GlcNAc, the supernatants were combined, and processed for sample preparation for mass spectrometry as given below.

Buffers for sample preparation for mass spectrometry

50 mM Ammonium bicarbonate (AMBIC) buffer: Dissolve 3.95 mg of ammonium bicarbonate powder in 1.0 mL of water (mass spectrometry grade) in a clean glass vial; vortex mix thoroughly before use.

200 mM Dithiothreitol (DTT) buffer: Dissolve 30.85 mg of DTT in 1.0 mL of AMBIC buffer in a clean glass vial; vortex mix thoroughly before use.

20 mM Iodoacetamide (IAA): Dissolve 3.7 mg of IAA in 1.0 mL of AMBIC buffer in a clean glass vial; vortex mix thoroughly before use.

### Sample preparation for mass spectrometry and bottom-up protein identification

Proteins (200 mL each from WT and KO samples, in 1.5 mL microcentrifuge tubes (Eppendorf)), eluted by GlcNAc from the lectin-affinity enrichment, were treated with ice-cold acetone (900 mL) and allowed to precipitated at −20 °C for 24 h. Samples were centrifuged (16000×g, 4 °C, 10 min) and the supernatants were aspirated and discarded. Protein precipitates were re-suspended in AMBIC buffer (100 mL), heated at 65 C for 10 min (at a shaking speed setting of 5 on a Torrey-Pines shaker). Samples were treated with DTT (2.5 mL; 200 mM) and heated at 60 °C for 45 min, followed by treatment with IAA (10 mL; 20 mM) at 37 °C for 45 min, on a shaker. Another aliquot of DTT (2.5 mL; 200 mM) was added to the samples and incubated for at 37 °C for 30 min on a shaker (Pv et al., 2024). Samples were treated with mass spectrometry grade trypsin (4.0 mL, 2.0 mg, Sequencing grade modified trypsin, Porcine, Promega, Cat. No. V511A) and incubated overnight at 37 °C. Samples were then treated with formic acid (1.0 mL) and concentrated to dryness in a refrigerated vacuum centrifuge at 4 °C. The samples were re-suspended in 5.0 % v/v aqueous acetonitrile containing 0.5 % v/v formic acid (30 mL) and de-salting was performed using reverse phase C-18 zip tips as given below.

### Desalting of tryptic digests using C-18 zip tips

1. First, acetonitrile (ACN) (10 mL) was used to wet the C-18 zip tips by aspiration using a P10 micropipette
2. Tips were washed using mass spec grade water with 0.1 % formic acid (FA) (10 mL) through aspiration by pipetting.
3. Samples (10 mL) were loaded on the zip tips through aspiration
4. Sample-loaded zip tips were washed using mass spec grade water with 0.1 % formic acid (FA) (10 mL × 2)
5. Samples were eluted using 50 % aq. ACN with 0.1 % FA (10 mL)
6. Samples were eluted for a second time using ACN (10 mL) and pooled in the same tube as in step 5.
7. Samples were evaporated to dryness in a refrigerated vacuum centrifuge at 4 °C.
8. Samples were re-suspended in 5% v/v aq. ACN with 0.1 % FA (12 mL).
9. Samples were subjected to nano-liquid chromatography tandem mass spectrometry (nano-LC-MS/MS) analysis at the NII Central Mass Spectrometry Facility (CMSF).

### Data acquisition using nano-LC-MS/MS, data processing, and data analysis

Desalted tryptic digests (5.0 mL per run) were loaded onto Vanquish Neo nano-LC (fitted with PepMap Neo C18 (5.0 mm, 300 mm × 5.0 mm, 1500 bar, trap/guard column and DNV PepMap Neo C18 (2.0 mm, 100 Å, 75 mm × 150 mm) capillary column) connected to an Orbitrap Exploris 240 High-resolution Mass Spectrometer (Thermo Fisher Scientific), equipped with a Nanospray Flex ESI source connected through an Nanobore stainless stell emitter (40 mm length; outer diameter 1/32 inch). Mobile phase A was kept as 0.1 % v/v formic acid in water and mobile phase B was kept as 0.1 % v/v formic acid in 80 % aqueous acetonitrile. Elution was performed using a multi-step gradient of 1.0 % to 99 % of mobile phase B from 0-100 min at a flow rate of 300 nL/min. Mass spectrometry data collection was performed in positive ion mode with electrospray at 1.9 kV, RF lens value at 70%, and ion-spray (1.9 kV) kept at 275 °C. MS1 scans were performed at a range of 375-1200 m/z with orbitrap resolution set to 60,000 and MS/MS scans at the resolution of 14000 in a data-dependent acquisition (DDA) mode. Higher-energy collision dissociation (HCD) was applied at a normalised collision energy (NCE) of 30%. Peptide with charge states of +2 to +6 were selected for fragmentation in MS/MS mode.

Raw data from nano-LC-MS/MS was processed using the XCalibur proteome discoverer (PD 3.0) software with SEQUEST HT as search engine. The acquired data was searched against UniProtKB complete mouse proteome database. Oxidation of Met was set as dynamic modifications and carbamidomethylation of Cys was set as a fixed modification. Parameters were set to a precursor mass tolerance of 10 ppm, fragment mass tolerance of 0.02 Da, and two missed cleavages allowed for trypsin. Proteins and peptide lists were compiled and represented as Venn diagrams using the protein distribution option of PD 3.0; Complete list for protein ID was generated separately for intra-samples (run to run). A list of probable target candidates of mgat4B were identified based on abundance ratio between control and KO samples. The proteins listed in the table were only present in wildtype samples and were absent in Mgat4b KO samples.

For label-free quantification, the processing was done by combining the technical runs for each sample. Label-free quantification (LFQ) was applied to identify relative quantitation of protein in control and KO samples. Label free quantitation was performed based on intensity of the precursor ion. For data processing, protein database was imported in the FASTA file format. The study factors were defined as knockout and wild type in the new study following which the raw data files were added to the study in .raw file format. The technical replicates of wildtype samples were defined as control and the knockout as sample. Under sequest HT node, enzyme selected was trypsin (full) with maximum missed cleavage sites of two, minimum peptide length of six amino acids (aa), maximum peptide length 144 aa, precursor mass tolerance 10 ppm, and fragment mass tolerance of 0.6 Da. Dynamic modification for oxidation (+15.995 Da) at methionine residue and static modification for carbamidomethyl (+57.021 Da) at Cystine residue were defined. Acetylation (+42.011 Da), methionine loss (-131.040 Da) and methionine loss with acetylation (-83.030 Da) at N-terminus were also selected under dynamic modifications (protein terminus). Dynamic modifications are the modifications which may or may not be present while the static modification is applied universally to every instance of the specified residues or terminus.

### Plasmid transfections in B16 mouse melanoma cells

B16 cells were trypsinized and seeded at density of 1 × 10^5^ cells/well in 6 well plate (nunc) and incubated overnight in antibiotic containing DMEM + 10% FBS. At the time of transfection, the cells were replaced with serum and antibiotic free media OptiMEM (Gibco, Life Technologies). Lipofectamine 2000 was used at a ratio of 1:3 with pcDNA 3.1(empty vector control) plasmid, Mgat4a and Mgat4b coding sequence containing plasmids. The cells were incubated for 6 hours with the transfection mixture containing Lipofectamine 2000, OptiMEM, and the plasmids. The media was then replaced with antibiotic containing DMEM + 10% FBS and incubated for 72 h, and downstream experiments were performed.

### Drug Treatments

Kifunensine (Cas no. 109944-15-2) was dissolved in DMSO and a final concentration of 150 µM was prepared in Embryo water. One-month old Mazerati zebrafish were treated for 2 days with a change of drug on each day.

### Imaging and Quantitative Analysis

Images of tumor area was captured using a Nikon SMZ800N camera. Brightfield images were adjusted for contrast and color balance for clarity. To quantify the effect of drug, tumor area was outlined and compared before and after treatment using ImageJ. Percent change in tumor area was calculated as a ratio of the change in tumor area across 2 days to the area covered by tumor after 2^nd^ day of kifunensine treatment. Student’s t tests were performed using GraphPad Prism 9.

### Molecular Docking

For MGAT4A, MGAT4B, MGAT4C and MGAT5 structure, Alpha-fold model (Varadi et al., 2022)was used since there were incomplete crystal strutures. The Model was pre-processed, polar hydrogen bonds added, kollman charges added and pdbqt structure was generated using Autodocktools. Grid box was defined for the models using Autodocktools. The ligand library of 2800 FDA approved drugs were energy minimized and converted to pdbqtformat using PyRx tool. Autodock Vina was used to do the virtual screening (Trott & Olson, 2010).

### Statistical analysis and graphs

Student’s t-test was performed to obtain statistical significance in the data. Asterisk on the error bar corresponds to *P ≤ 0.05, **P ≤ 0.01, ***P ≤ 0.001, ****P ≤ 0.0001 and ns P > 0.05. Graphs were plotted using GraphPad prism.

## Ethics statement

Fish experiments were performed in strict accordance with the institutional animal ethics approval (IAEC) of the CSIR-Institute of Genomics and Integrative Biology (IGIB), India (Proposal No 45a). IEAC approval number–IGIB/IAEC/25/28/2020. All efforts were made to minimize animal suffering.

---

**Supplementary Figure 1:**
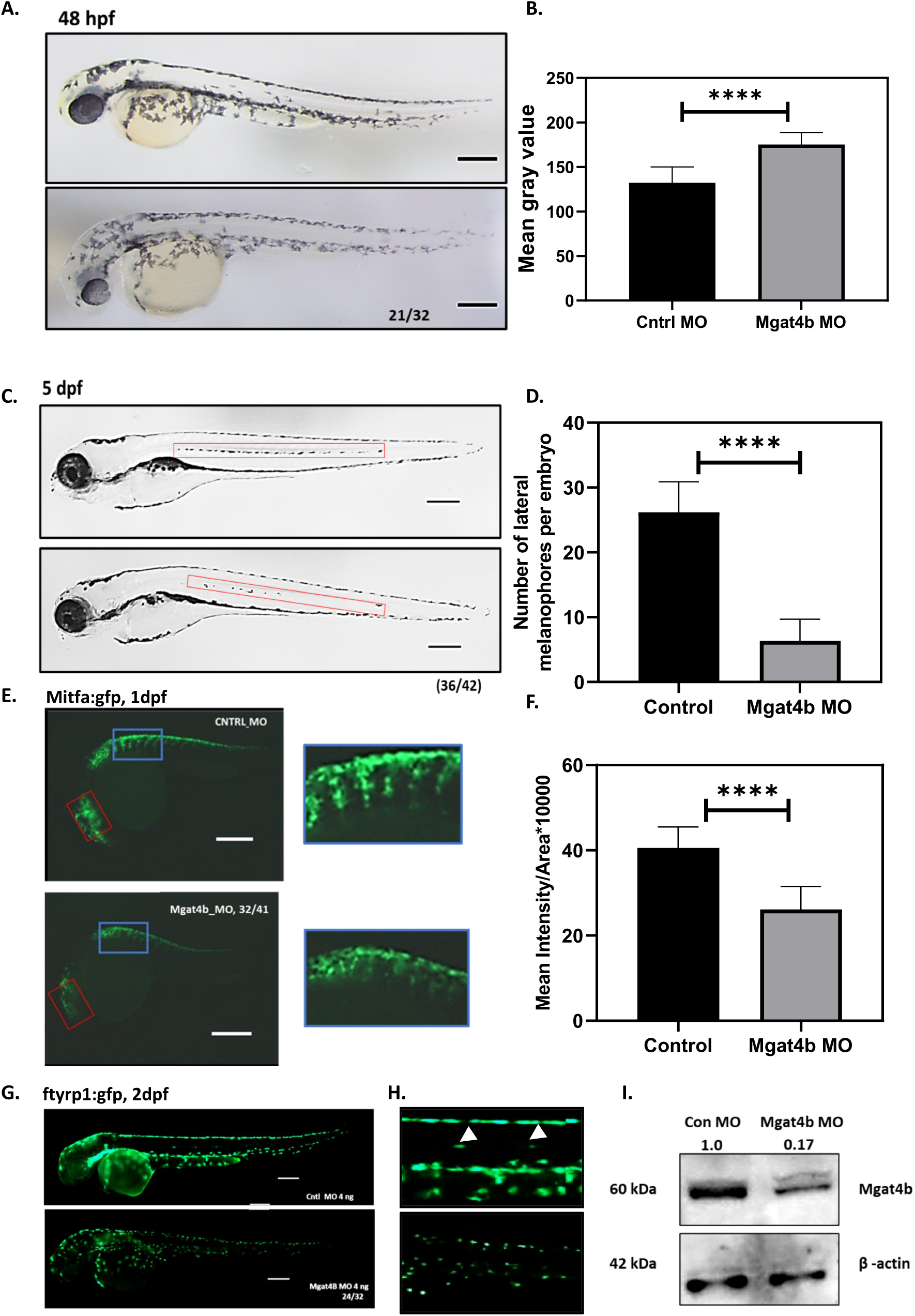
Targeting *mgat4b* during early zebrafish development leads to pigmentation and melanophore patterning defects: **A**. Brightfield images of the lateral view of control and *mgat4b* morphant embryos at 2 days post fertilization (dpf). The black structures observed are melanophores, Scale bar-100µm **B**. Melanin quantitation from the brightfield images of control and *mgat4b* morphants was carried out using ImageJ platform. The mean gray values are inversely linked to melanin content of the embryo and are represented as a Bar graph **C**. Brightfield images of the lateral view of control and *mgat4b* morphant embryos at 5dpf, Scale bar-100 µm **D**. The number of lateral mid-line melanophores were counted manually (highlighted in red box) represented in the graph. Scale bar-100µm **E.** Fluorescent images of control and *mgat4b* morphants at 1 dpf. Red boxes mark the constant region chosen to quantify mean fluorescent GFP intensity per embryo, Scale bar-100 µm **F.** Mean intensity per area is quantified and depicted in the bar graph **G.** Fluorescent images of control and *mgat4b* morphants at 2 dpf, Scale bar-100 µm **H.** Zoomed inset of 2dpf ftyrp:gfp control and *mgat4b* morphant trunk region. Scale bar-100µm G. Western blot analysis shows Mgat4b protein levels in bulk (100 embryos) of control and *mgat4b* morphants at 2dpf *P ≤ 0.05, **P ≤ 0.01, ***P ≤ 0.001, ****P ≤ 0.0001 and ns P > 0.05

**Supplementary Figure 2.**
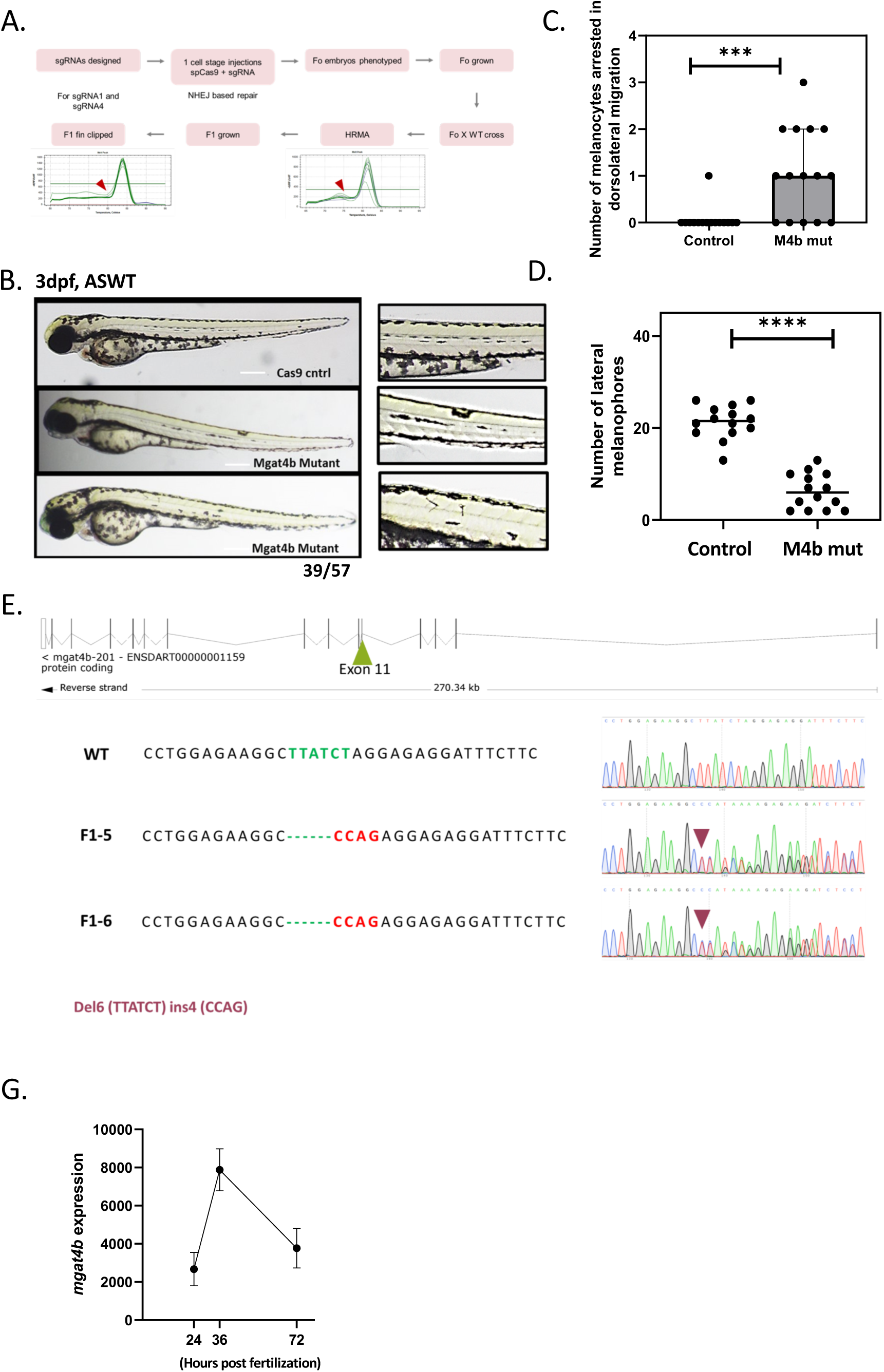
Global ablation of *mgat4b* shows patterning defects. **A**. Schematic representation of workflow acquired to create global mgat4b *mutant* in zebrafish **B.** Brightfield images of F0 cas9 control and *mgat4b* mutant at 3dpf stage **C.** Bar plot depicting the number of melanophores arrested in between the path **D.** Bar plot depicting the number of lateral melanophores at 3 dpf stage of F0 cas9 control and *mgat4b* mutant Scale Bar 100um **E.** Validation of F1 *mgat4b* knockout animals by sanger sequencing **F.** Dot plot depicting the RNA levels of *mgat4b* in melanophores at three different stages of their development (24, 36 and 72 hours post fertilization). Re-analysed Zebrafish time-course microarray data (GSE189059) *P ≤ 0.05, **P ≤ 0.01, ***P ≤ 0.001, ****P ≤ 0.0001 and ns P > 0.05

**Supplementary Figure 3.**
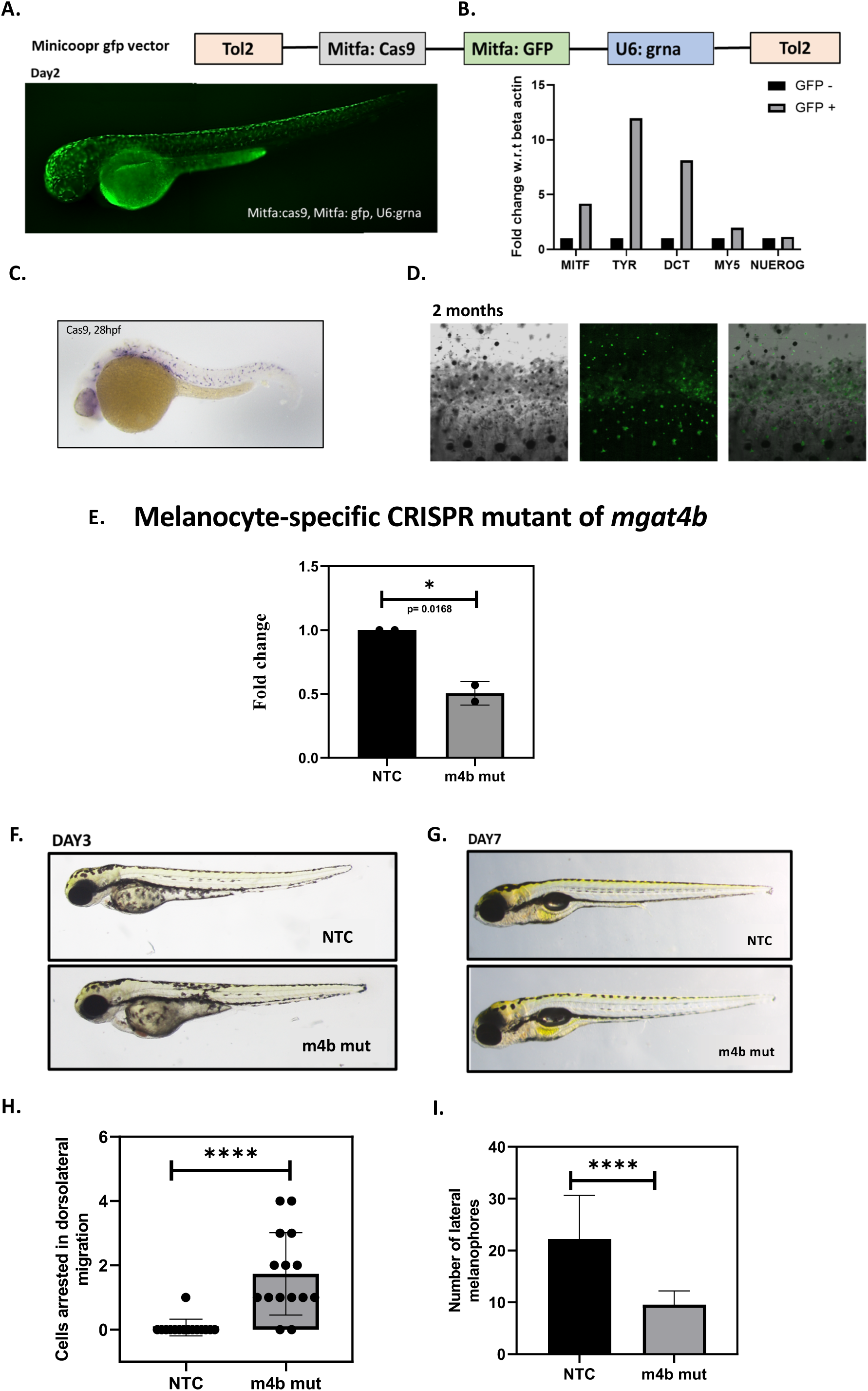
Melanocyte-specific ablation of genes and visualization in zebrafish. **A.** Fluorescent image of 2dpf injected with mitfa:cas9;mitfa:gfp plasmid at a single cell stage **B.** Bar plot showing transcriptomic levels of melanocyte specific genes mitf, tyr and dct in FACS sorted GFP+ cells, muscle and neuron related negative controls are also shown **C.** WISH shows Cas9 expression in 28hpf zebrafish injected with mitfa:cas9;mitfa:gfp plasmid at a single cell stage **D.** Confocal images of 2 month old zebrafish injected with mitfa:cas9;mitfa:gfp plasmid at a single cell stage shows gfp+ cells **E.** qRT-PCR for *mgat4b* transcripts in FACS sorted gfp+ cells upon tissue-specific targeting of *mgat4b* in melanophores. Fold change is depicted (mean ± SEM, n ≥ 2, biological replicates) calculated using beta actin as the reference with respect to non-targeting control animals **F.** Brightfield lateral images of tissue-specific knockout of *mgat4b* (m4b mut) and NTC (Non-targeting control) animal at 3dpf **G.** Brightfield lateral images depicting melanophore stripes of m4b mut and NTC animal at 7 dpf **H.** Bar plot depicting the number of melanophores arrested in between the path for m4b mut and NTC animals **I.** Bar plot depicting the number of lateral melanophores in m4b mut and NTC animals *P ≤ 0.05, **P ≤ 0.01, ***P ≤ 0.001, ****P ≤ 0.0001 and ns P > 0.05

**Supplementary Figure 4.**
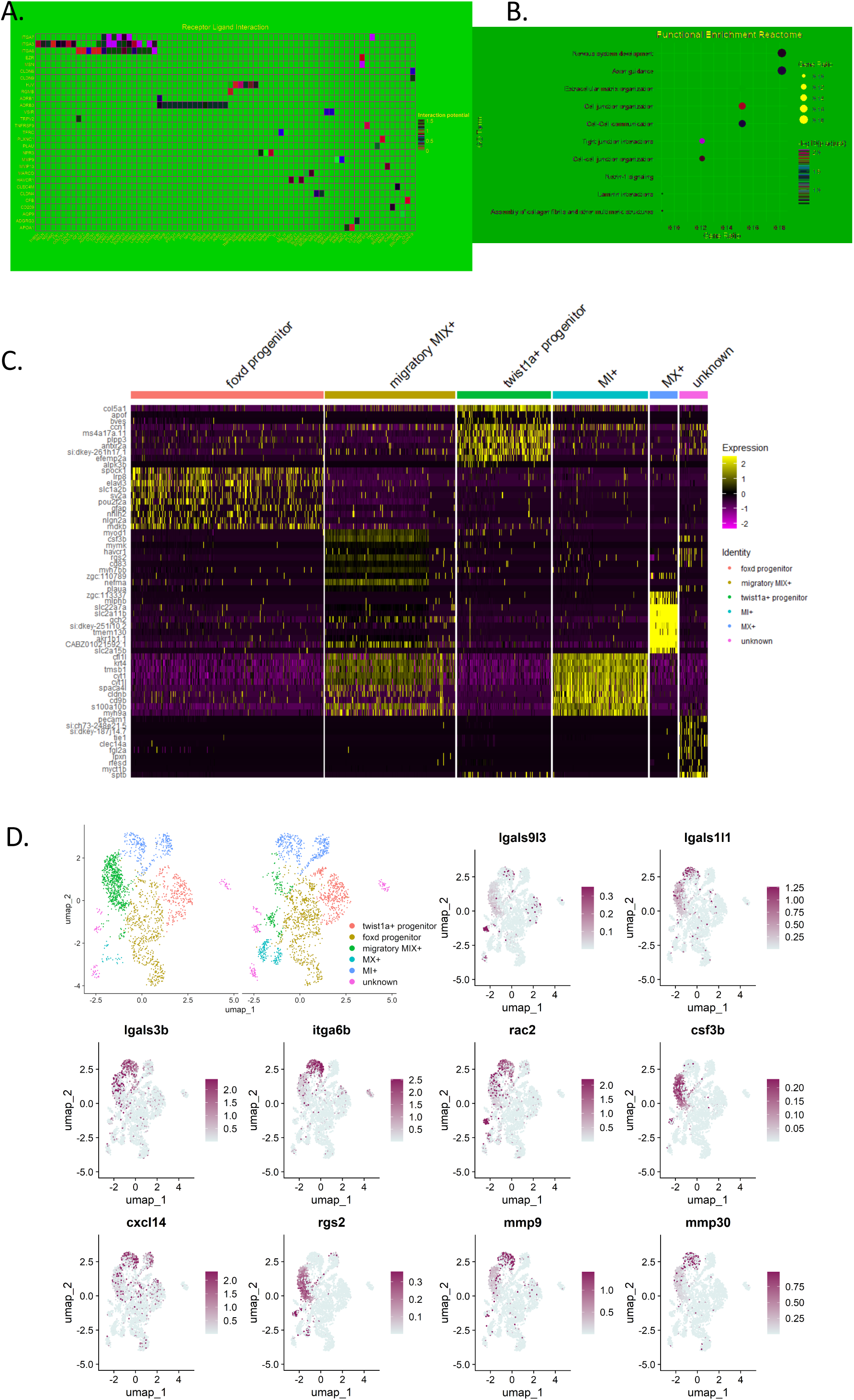
scRNA seq reveals the identity of the depleted melanocyte population. **A.** Heatmap of NicheNetR-identified ligand/receptor pairs indicating interaction potential between MIX+cluster cells and other sender cells sampled by scRNAseq **B.** Functional enrichment for reactome of top enriched marker genes of MIX+ cluster **C.** Heat map depicting top markers enriched in each cluster and used to annotate them **D.** UMAPs of Mitfa+ cells in NTC and m4b mut with color change from gray (negative) to purple based on log normalized scaled expression of lgals9I3, lgals91I1, lgals3b, itga6b, rac2, csf3b, cxcl14, rgs2, mmp9, mm30 genes

**Supplementary Figure 5:**
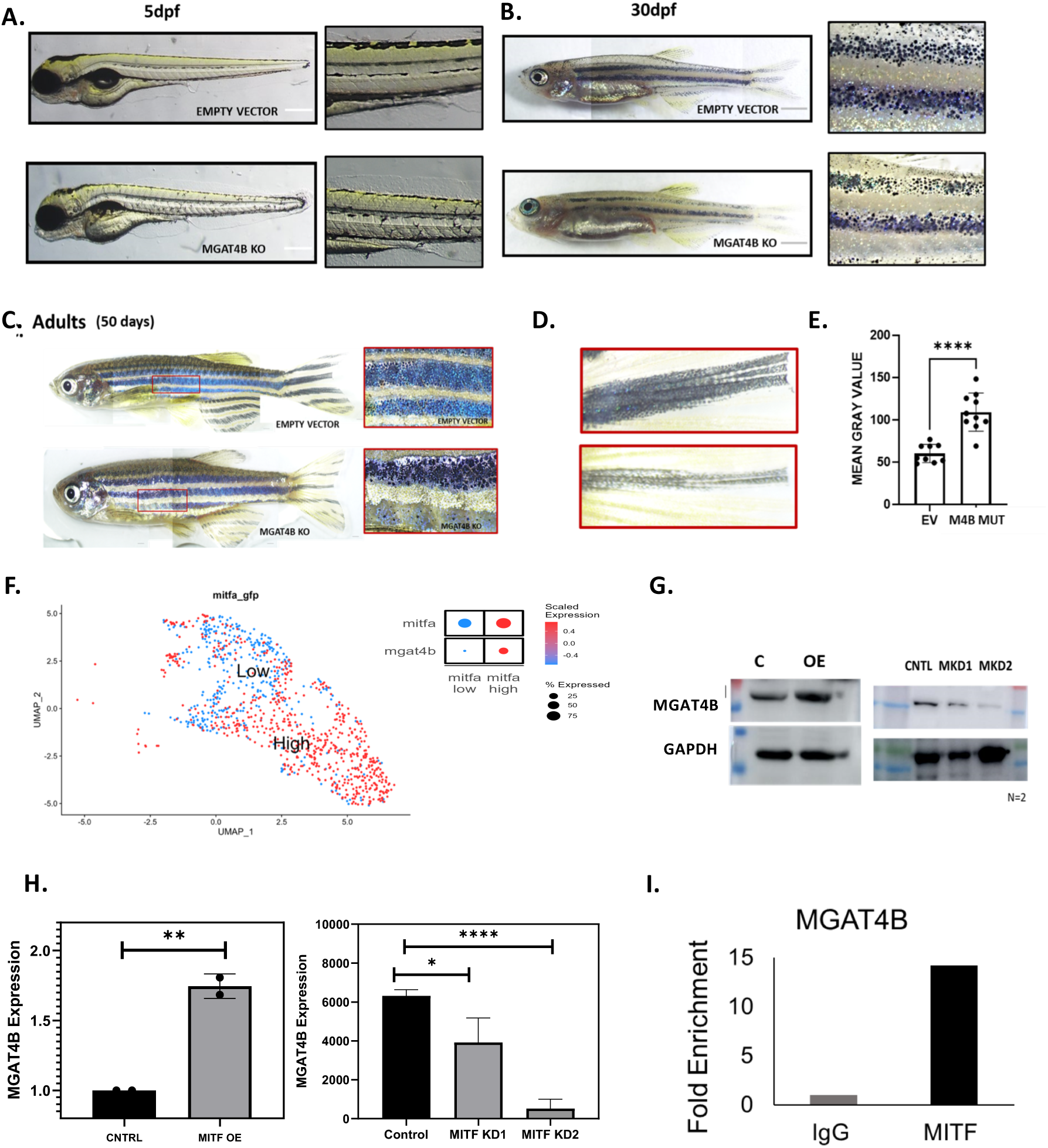
Central transcription factor MITFA transcriptionally regulates levels of MGAT4B in melanocytes: **A.** Lateral images of 5dpf NTC and melanocyte specific *mgat4b* mutant (m4b mut) animals showing embryonic melanocytic stripes **B.** Lateral images of 30dpf NTC and m4b mut animals showing adult stripe patterns, zoomed images shows pigment stripes in the trunk region **C.** Lateral images of 50dpf NTC and m4b mut animals showing adult stripe patterns, zoomed images shows piment stripes in the trunk region **D**. Zoomed images of tail fin from 50dpf NTC and m4b mut animals **E**. Bar graph represents mean±s.e.m. of mean gray value of tail fin selected ROI in NTC and m4b mut, each dot represents an animal **F**. UMAP plot showing Mitfa high (red) and Mitfa low(blue) cell clusters, dot plot showing collective expression of Mitfa and mgat4b in these cells **G.** Western blot showing levels of MGAT4B upon overexpression of MITF in B16 mouse melanoma cells, Western blot showing expression of MGAT4B upon downregulation of MITF using two different siRNAs **H)** Bar graph showing quantitation of D and E **I.** Bar plot showing fold enrichment of MITF over MGAT4B promotor region revealed by ChIP-qPCR *P ≤ 0.05, **P ≤ 0.01, ***P ≤ 0.001, ****P ≤ 0.0001 and ns P > 0.05

**Supplementary Figure 6:**
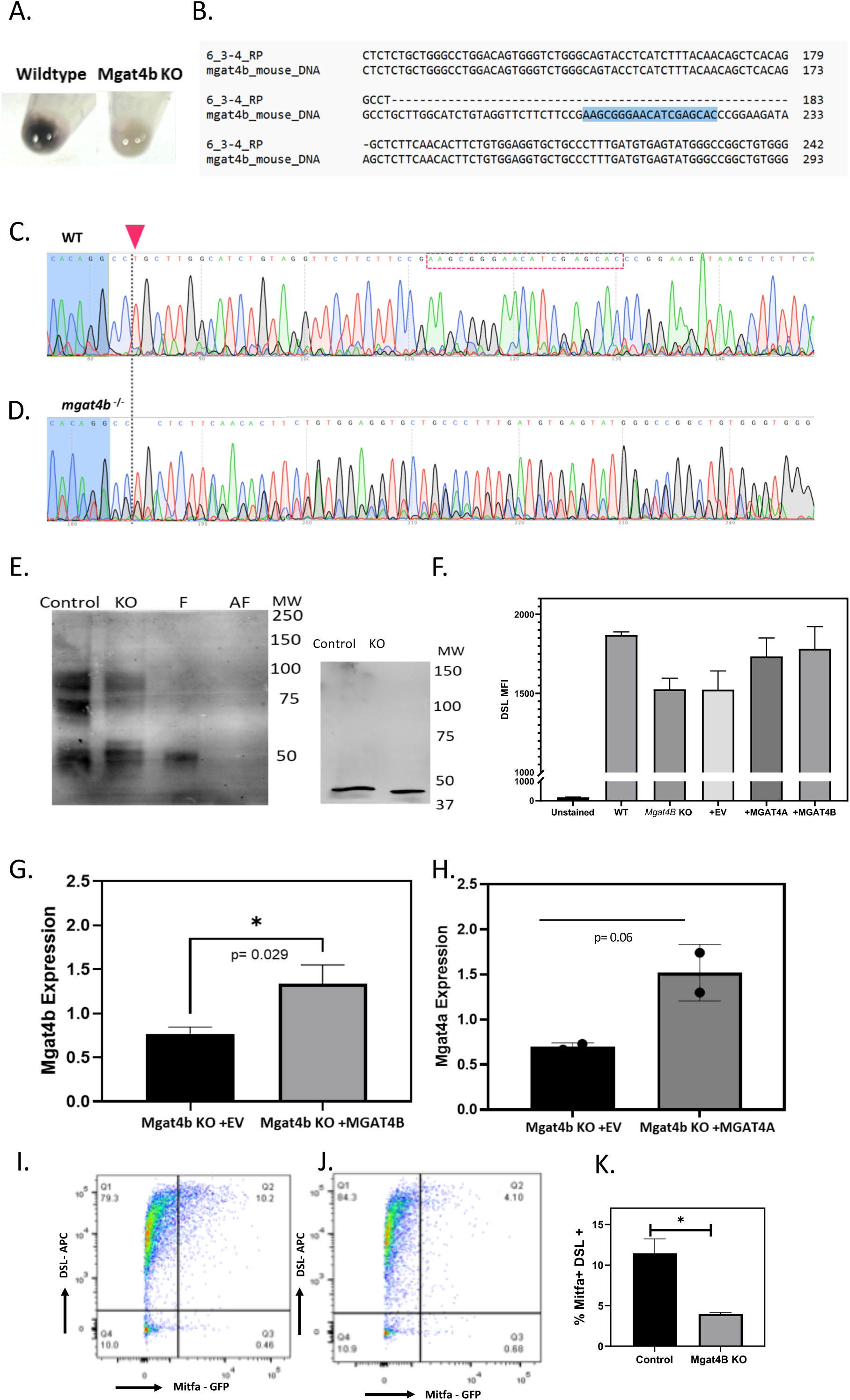
***Mgat4b* knockout leads to change in melanoma cells properties. A.** Cell pellet of day 7 B16 mouse melanoma wildtype and *Mgat4b* KO cells grown at low density (100 cells/cm2) **B.** Sanger sequencing based validation of *Mgat4b* knockout (colony 6, Mut1) in B16 mouse melanoma cells compared with wildtype sequence **C, D.** Sanger sequencing chromatogram of wild type (WT) and *Mgat4b* knockout (*Mgat4b*^-/-^) clone shows deletion of 57 bp around the highlighted sgRNA region **E.** DSL lectin blot depicting enrichment in wildtype and *Mgat4b* KO whole cell lysate with positive control (Fetuin) and negative control (asialofetuin) in place, beta-actin levels assessed as a loading control **F.** Bar plot shows mean±s.e.m of DSL lectin enrichment on the cell surfaces of Wildtype, *Mgat4b* KO, *Mgat4b* KO cells complemented with Empty vector, MGAT4B and MGAT4A protein **G, H.** Mgat4b and Mgat4a expression in Mgat4b KO cells upon complementation of MGAT4A and MGAT4B protein fold change calculated wrt B16 WT cells **I, J.** FACS plot shows enrichment of DSL lectin on the cell surfaces of zebrafish NTC and *mgat4b* mutant cells **K.** Bar plot depicting the frequency of double positive cells (GFP+, DSL+) in NTC and *mgat4b* mutant melanocytes *P ≤ 0.05, **P ≤ 0.01, ***P ≤ 0.001, ****P ≤ 0.0001 and ns P > 0.05

**Supplementary figure 7:**
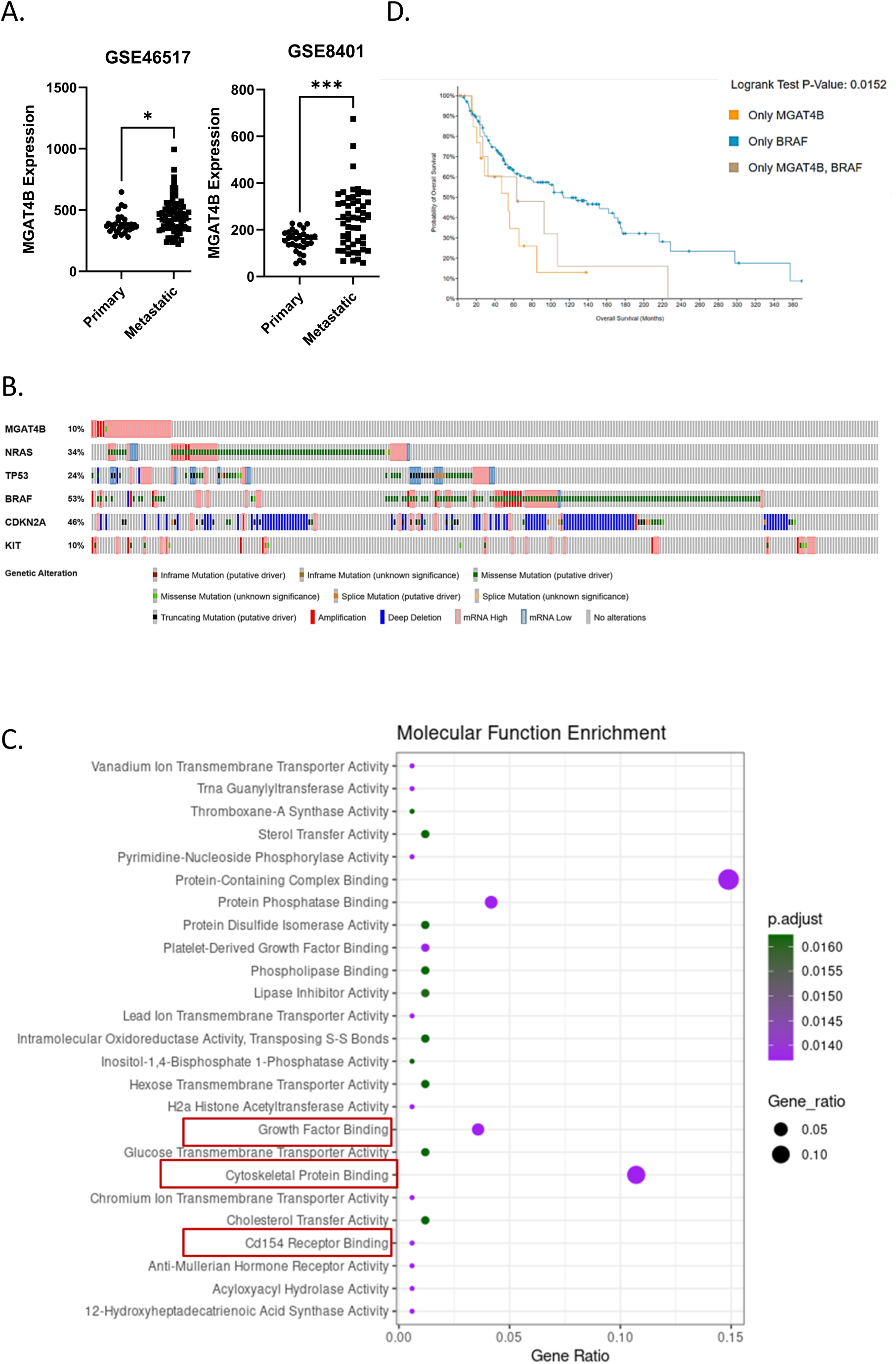
Re-analysis of publicly available data shows enrichment of MGAT4B in melanoma patients. **A.** Scatter plot depicts transcriptomic levels of MGAT4B in skin cutaneous melanoma patients segregated based on primary or metastatic stage, two different datasets were meta-analysed with the GSEA ID-GSE8401, GSE46517 **B.** Alteration frequency of candidate genes in the TCGA melanoma cohort, Top altered genes are plotted to display deep deletions, missense mutations, and amplifications along with MGAT4B **C.** Molecular function enrichment of differentially regulated genes between MGAT4B high and MGAT4B low SKCM patients, data procured from TCGA **D.** Kaplan–Meier survival plots comparing overall survival (OS) probabilities (Y-axis) as a function of time in months (*x*-axis) in melanoma patients with mutated BRAF, MGAT4B and both BRAF and MGAT4B p value 0.0152

**Supplementary figure 8:**
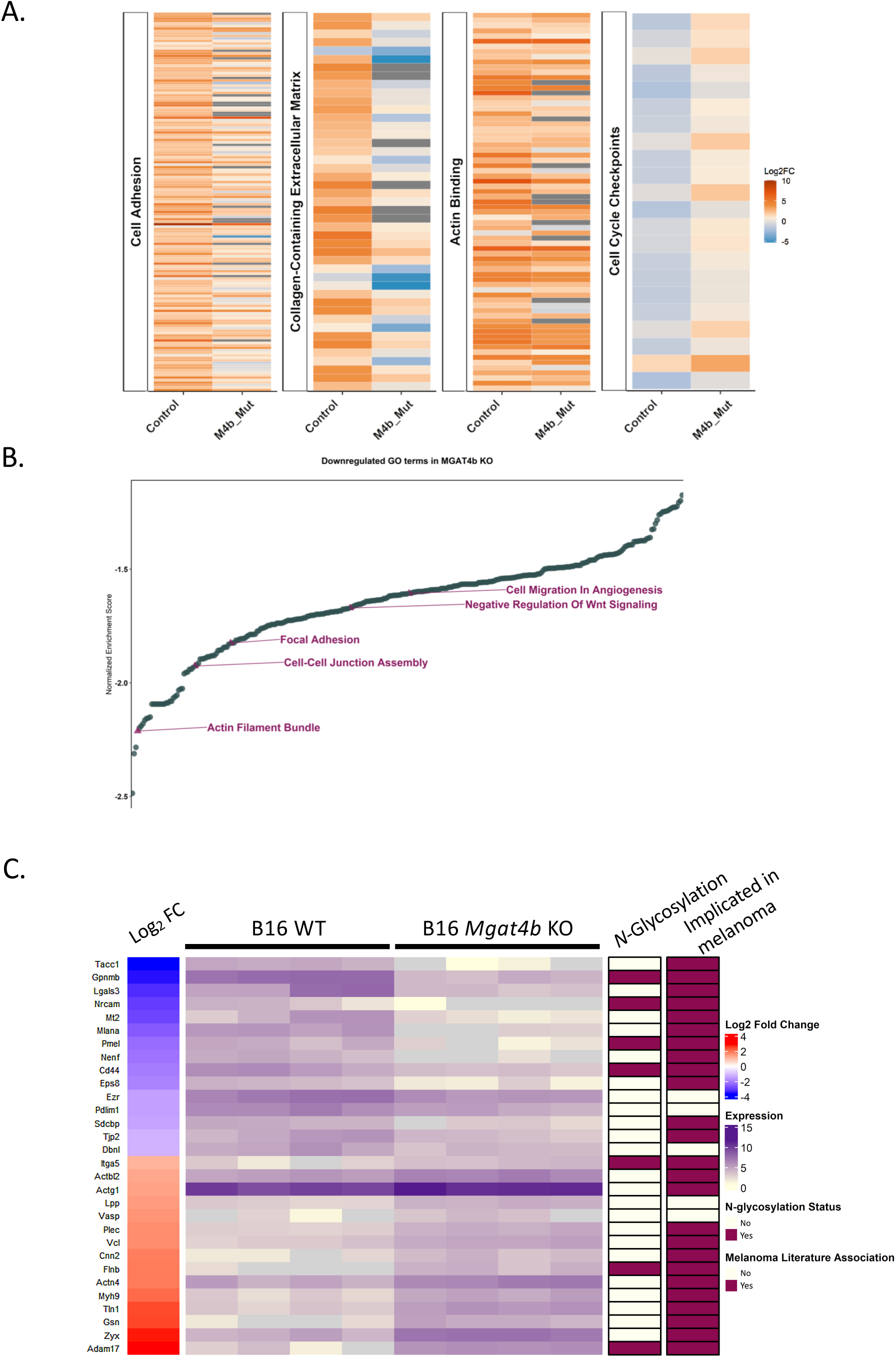
Altered Cell Adhesion in zebrafish *mgat4b* mutants and B16 KO Cells: **A.** Heap map depicting log2 fold change of genes related to the GO terms cell adhesion, collage-containing extracellular matrix, actin binding and cell cycle check points in control melanoma (control), *mgat4b* mutant (M4b_Mut) melanoma with respect to wildtype melanophores **B.** Waterfall plot depicting normalized enrichment score of downregulated GO terms in *Mgat4b* KO with respect to wildtype cells (B16 mouse melanoma), keys terms are highlighted **C.** Heap map depicting log_2_ fold change, log_2_ expression, *N*-glycosylation status and melanoma association of top differentially regulated proteins from B16 WT and B16 *Mgat4b* KO cells.

**Supplementary Figure 9:**
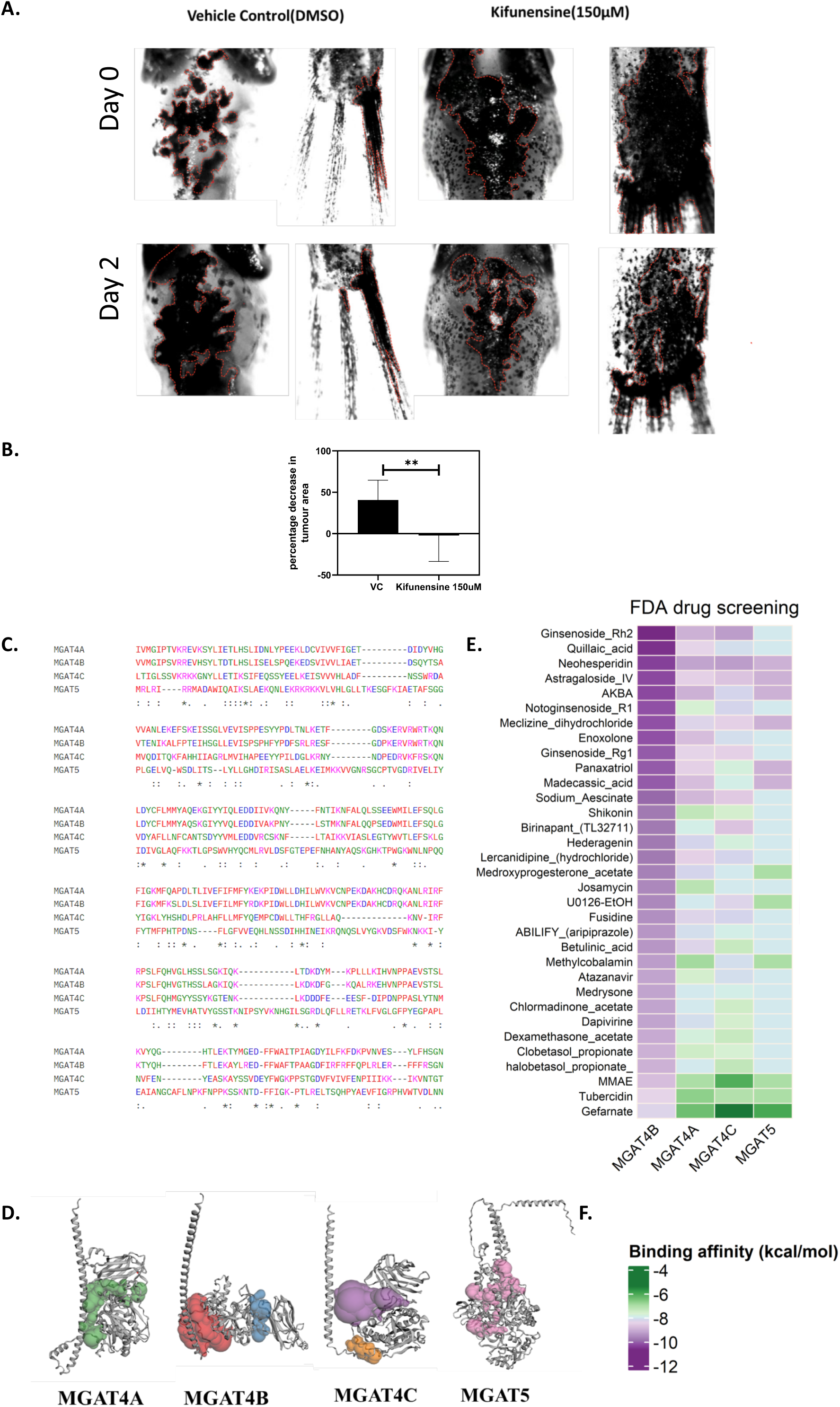
Inhibition of complex *N*-glycosylation as a potential therapeutic intervention for melanoma initiation and progression. **A.** One-month-old zebrafish with genetically initiated melanoma tumors, as previously described, were treated with the drug kifunensine. Grayscale images show the tumors at day 0, prior to treatment, and at day 2, after two days of drug treatment (150µM) in both the treated group and the vehicle control (DMSO) group **B.** Bar plot depicting mean±s.e.m of the percentage decrease in tumor area upon kifunensine treatment and vehicle control group **C.** Multiple Sequence Alignment of the protein sequences pertaining to Human MGAT4A, MGAT4B, MGAT4C AND MGAT5 **D.** alpha-fold models of MGAT4A, MGAT4B, MGAT4C AND MGAT5 with their predicted binding pockets highlighted **E, F.** Heat map depicting the binding affinities of MGAT4A, MGAT4B, MGAT4C AND MGAT5 with the prioritised drugs. *P ≤ 0.05, **P ≤ 0.01, ***P ≤ 0.001, ****P ≤ 0.0001 and ns P > 0.05

**Supplementary Table 1**: Cells velocity in the chemotaxis chamber when provided with a chemoattractant (SCF) added media or neutral media to control and B16 Wild type and B16 Mgat4b KO cells

**Supplementary Table 2**: Proteomic analysis of whole cell lysates from wild type and Mgat4b KO B16 melanoma cells

**Supplementary Table 3**: Proteomic analysis upon enrichment by DSL lectin in whole cell lysates of B16 Wild type and B16 Mgat4b KO cells

**Supplementary Table 4**: Transcriptomic analysis of control and *mgat4b* mutant melanoma/melanophores from MAZERATI fishes with respect to wildtype melanophores from stage matched zebrafish

**Supplementary Table 5**: List of markers utilized for annotating the specific clusters for the scRNA seq data

**Supplementary Table 6**: List of FDA approved drugs along with their predicted binding affinities to MGAT4A, MGAT4B, MGAT4C and MGAT5.

**Supplementary Video 1**: Time lapse imaging of 28 hpf zebrafish injected with Mitfa:cas9;Mitfa:GFP: U6 sgRNA non-targetting plasmid as a control group

**Supplementary Video 2**: Time lapse imaging of 28 hpf zebrafish injected with Mitfa:cas9;Mitfa:GFP: U6 *mgat4b* sgRNA plasmid as test group

## Data Availability statement

Transcriptomic data pertaining to single cell sequencing and RNA sequencing has been submitted to GEO repository

**GSE278653** Mgat4b mediated selective *N*-glycosyl modification regulates melanocyte development and melanoma progression [bulk RNA-seq]

**GSE278654** Mgat4b mediated selective *N*-glycosyl modification regulates melanocyte development and melanoma progression [scRNA-seq]

Proteomic data pertaining to Lectin Affinity enrichment and whole cell expression from B16 WT and B16 *Mgat4b* KO cells submitted to ProteomeXchange Consortium via the PRIDE partner repository

**PXD056636** Proteomic data from B16 WT and B16 *Mgat4b* KO whole cell lysate

**PXD056675** Proteomic data from Lectin affinity enrichment using DSL from B16 WT and B16 *Mgat4b* KO whole cell lysate

## Acknowledgement

We thank Akshay Anil Binayke, Kritika Sahoo, Dharini Alagar Selvam, Shivani Chitkara and Manish Chowdhury from IGIB for help with experiments and analysis. We thank Richa Arya and Deepanshu Sharma, Central Mass Spectrometry Facility (CMSF), NII, for data acquisition and data processing on differential proteomics study of lectin-affinity enriched samples. We thank Prof Ramray Bhat from Department of Developmental Biology and Genetics Indian Institute of Science and Dr Sridhar Sivasubbu from CSIR-IGIB for useful discussions.

